# A big and hairy business: Investigating the interplay of species traits and trade dynamics in the tarantula pet market

**DOI:** 10.1101/2023.07.28.550999

**Authors:** Caroline S. Fukushima, Adam Toomes, Diogo Veríssimo, Pedro Cardoso

## Abstract

Species traits significantly influence pet trade dynamics, affecting demand, exploitation, and extinction risk. We examined the effect of species- and advertisement-level attributes on tarantula abundance and price in online markets, exploring rarely-considered fine-scale traits. Data from 977 ads showing 217 species and 81 ‘trade names’’ were collected from eight e-commerce websites located in six countries and analyzed using Structural Equation Models. Hairy, aggressive, and popular tarantulas were more abundant in commerce. Big, recently described species with ontogenetic changes in color and urticating hairs, with no evidence of captive breeding, had higher average price. Variability of prices in the ads were mainly explained by differences in website, lifestage and sex of the individual advertised. After accounting for these drivers of market abundance and price, we found only weak evidence of direct price-rarity relationships, implying they are largely independently determined. This can have important implications for the future management and regulation of the international tarantula trade. Understanding consumer behavior in the pet trade is crucial for effective conservation efforts and we recommend using online ad data to track and understand supply and demand in tarantula trade. Leveraging marketing insights can enhance conservationists’ influence on consumers, promoting sustainable practices and benefit sharing for wildlife-dependent communities. Hobbyist communities may be the most strategic messenger for conservation messaging trying to reach consumers in the tarantula pet trade.

**Article impact statement:** Tarantula price and abundance in pet trade are shaped by species traits and advertisement attributes.

## Introduction

The desire for exotic (wild or non-domesticated) pets has existed for at least the last millennium of human culture (Driscoll & Macdonald 2010), being driven by a diverse array of motivations (Hausmann et al. 2023). In the modern era, the exotic pet trade is associated with a host of risks including the spread of zoonotic diseases, transport and release of invasive alien species, and unsustainable harvest of threatened populations (Lockwood et al. 2019, Marshall et al. 2022). As such, there is a clear need to study pet trade dynamics, identify the taxa involved, and build predictive capacity to pre-emptively identify trade-based threats. However, the vast majority of contemporary research on pet trade dynamics and impacts focus on ‘charismatic’ vertebrate taxa (Bush et al., 2014; Fukushima et al. 2020). Many other animals, particularly invertebrates like mantids, beetles, ants, snails, millipedes, and arachnids are traded as pets (Losey et al. 2022, Battiston et al. 2022, Lassaline et al. 2023). Tarantulas (Araneae, Theraphosidae), for example, with their exotic appeal and low maintenance requirements, have gained popularity as pets since the 1970s, leading to the formation of several hobbyist tarantula societies worldwide. Since then, their trade has been growing, currently encompassing at least 50% of the 1000+ described species (Marshall et al. 2022).

### The role of species traits

Understanding the drivers of exotic pet trade is crucial for effective conservation strategies. Ecological and morphological traits play a pivotal role in shaping the dynamics of the trade, since they can influence species demand and consequently increase exploitation and extinction risk (Hughes et al. 2023). The identification of preferred traits can help in different ways. First, it aids to understand why particular species become targets (Battiston et al. 2022). Second, it provides valuable insights into related species with similar desirable features that can become targets when the abundance of the primarily targeted species declines or when they become known to prospective traders (Vall-llosera & Cassey 2017). Finally, identifying traits that drive market value and abundance may predict the capacity to anticipate biosecurity threats posed by trade (Vall-llosera & Cassey 2017). For instance, studies have demonstrated that species sold at lower prices are more frequently released compared to rare and expensive species (Stringham & Lockwood 2018; Macega-Veiga et al. 2019).

### Trait-trade dynamics

Trait-trade dynamics are typically studied at course taxonomic levels (e.g., Aves; Toomes et al. 2022) using traits that are widely available through databases (e.g., Amniote; Myhrvold et al. 2015). As more taxa-specific traits may be driving finer-scale patterns, exploring datasets of these traits can uncover nuanced patterns, ultimately enhancing our ability to make informed decisions regarding conservation and trade management strategies. For instance, traits such as the presence of complex mating and hunting behaviors were suggested as relevant factors in species purchase (Battiston et al. 2022). Novelty and uniqueness were also suggested to be key drivers in the trade of tarantulas (Marshall et al. 2022), but no quantitative analysis was performed to confirm those propositions.

### Monitoring online markets

The rise of the internet and online communities fostered further connectivity among tarantula enthusiasts through specialized forums and social media platforms. While the shift of trade to online marketplaces has facilitated a growth in the business (subsequently increasing the potential biosecurity and conservation risks), it simultaneously provides an opportunity for large-scale monitoring of trade. Such monitoring may help reveal consumer preferences for tarantulas, provide insights into the relationship between demand and price, and detect temporal changes and trends (Sung and Fong 2018).

Tarantulas represent a taxon that is (i) highly prevalent in trade, including online trade, yet poorly studied (Shivambu et al. 2020; Marshall et al. 2022); (ii) particularly susceptible to unsustainable harvesting due to slow life-cycle and relatively restricted species distribution (Marshall et al. 2022); (iii) a potential future source of invasive species via the pet-release pathway (Shivambu et al. 2020); and (iv) poorly represented in conservation agreements and policies (Fukushima et al. 2019). As such, it is necessary to investigate the international trade of tarantulas as exotic pets while considering traits which may make them specifically desirable to keepers and traders in the invertebrate pet hobby. Specifically, we intend to determine which traits at both species and advertisement level influence the abundance and price of specimens in online markets, providing a characterisation of trait-trade dynamics for fine-scale traits that are rarely considered.

## Material and Methods

### Online markets

Data was collected manually from e-commerce websites dedicated to tarantula sales in July 2018. In order to find potential candidates for sourcing the advertisements (ads hereafter), we utilized the Google search engine, employing Boolean search strings such as ‘tarantula AND store AND Germany’ (all terms in English) to identify e-commerce platforms. Similar searches were conducted for countries with established tarantula hobbies like the UK, USA, Germany (CEC 2017), as well as countries with large potential markets (sizable populations and/or presence of native tarantula species) such as China, Brazil, India, and Indonesia. As a result, we found either links to e-commerces or links to blogs, forums and other sites (e.g. Arachnoboards, Tarantula Forum) where people were referring to places selling tarantulas in the searched country. We aimed to ensure that the selected websites were more likely to represent establishments with a substantial inventory and a greater impact on the overall abundance and pricing of tarantula species in the online market. Thus we focused on e-commerces specialized in tarantula trade that displayed at least 30 advertisements in their page. The threshold of 30 advertisements was established after considering information gathered from prior searches, which revealed that many non-specialized and/or not established tarantula-selling places had less than 15 ads at the time of inspection. Additionally, we looked for websites with positive reviews and those that provided detailed information about care requirements, shipping policies, and guarantees, as these factors may indicate a reputable and established business.

### Taxonomy

We standardized scientific names according to The World Spider Catalog v.19.0 (2018), the version available at the time of data collection. We considered the names advertised as ‘operational taxonomic units’ (OTUs), regardless of their taxonomic validity, as we assumed that buyers were seeking specific names when purchasing a specimen and may not be aware of changes in tarantula taxonomy. Therefore, we did not aggregate synonyms. Furthermore, synonyms may differ from one another in various categories, such as year or description, global interest, or last publication’s authorship in the WSC, all variables of interest to our analysis. Establishing the identity of ‘trade names’ can often be challenging since many of them may represent undescribed species, hybrids, or the same name being used for different morphotypes depending on the place of sale (Toomes et al. 2023).

### Advertisements

We considered ads selling any number of individuals of any single taxon within Theraphosidae. ‘Out of stock’ ads were excluded since they do not represent a genuine possibility of purchase for the consumer. Bulk orders were excluded (except to calculate abundance per species) because certain attributes (e.g., sex, lifestage) may not be consistent across all individuals advertised in a bulk sale. Any duplicated adverts were excluded. Prices on the ads were given in different currencies but were converted to Euros based on the Euro exchange rate taken on 29 June 2018 (https://www.ecb.europa.eu/stats/policy_and_exchange_rates/euro_reference_exchange_rates/html/index.en.html).

### Species traits variables

We selected species-level traits that experts, including ourselves (CSF and PC), have hypothesized to be relevant in the tarantula pet trade (see Figure 1, Table 1 and Appendix S1 for mode details). In addition, we included variables that have been identified in the literature as potentially influencing trade volume or price in the wildlife trade for other groups, such as presence in the Appendix of the Convention on International Trade in Endangered Species of Wild Fauna and Flora (hereafter CITES) and in the Global Red List of the International Union for Conservation of Nature (hereafter IUCN), and traits related to husbandry (Vall-llosera and Cassey 2017). Categorical variables were treated as ordinal if there was reason to assume that categories were ordered in terms of their effect on the response (e.g., slow, medium & fast growth rate). Some species attributes that could be relevant for the trade dynamics, such as number of offspring, species longevity, and some behavioral traits were not included due to poor data availability and/or reliability.

**Figure 1.**
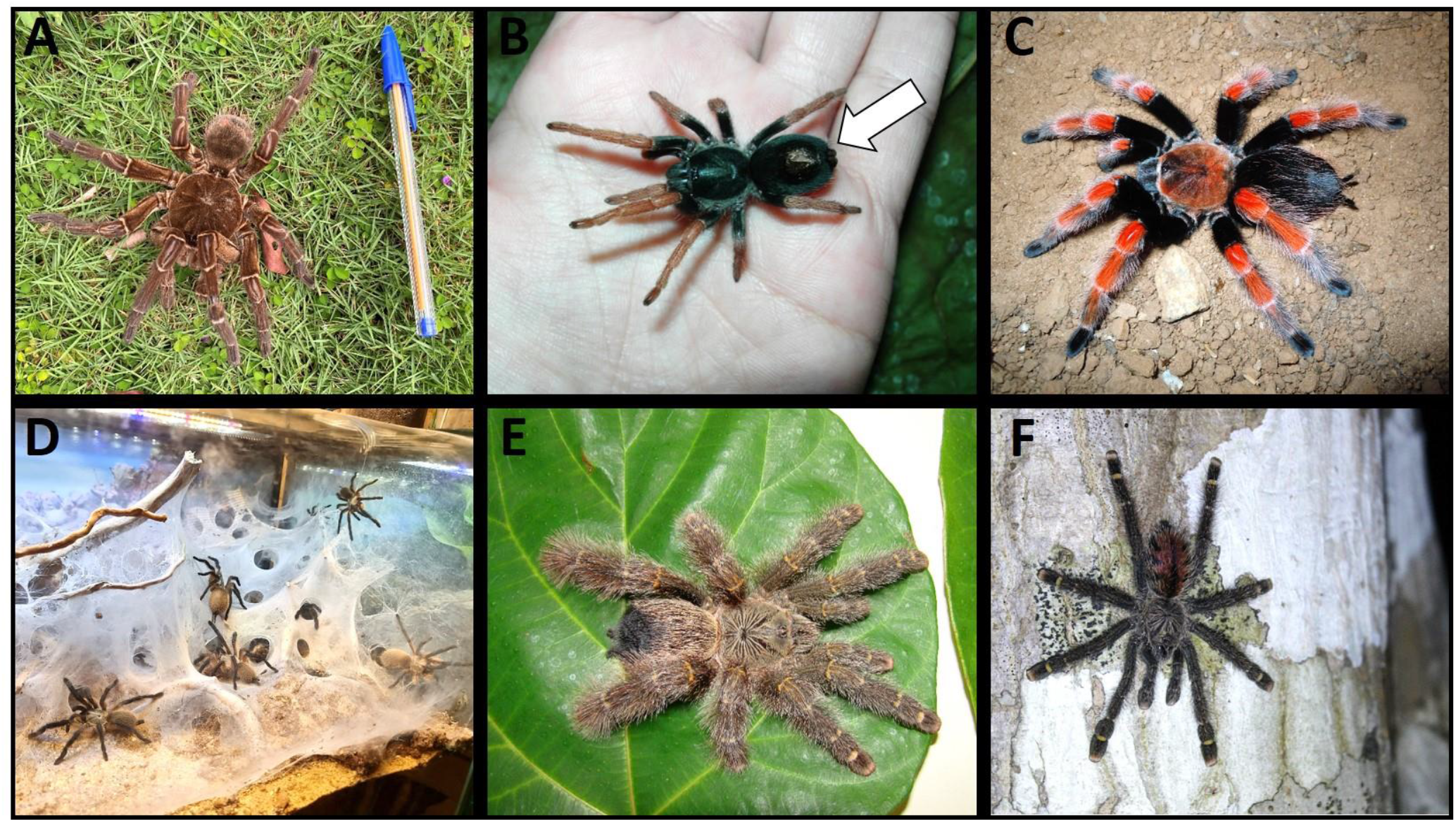
Tarantula species showcasing their corresponding traits utilized in the analysis, denoted within parentheses. A. Goliath birdeater *Theraphosa blondii* (Latreille, 1804) – Size (large category) and hairiness (low score). B. Dwarf Pink Leg *Kochiana brunnipes* (C.L. Koch, 1841) – Size (small), docility (docile), and urticating hairs (presence). Arrow indicates the patch of abdominal urticating hairs. C. Mexican Fire Leg *Brachypelma boehmei* Schmidt and Klaas, 1993 – Color (high score), Regulatory status (present in CITES), and Threatened status (evaluated in a threatened category in the IUCN Red List). D. Socotra Island Blue Baboon *Monocentropus balfouri* Pocock 1897 – Tolerance to conspecifics (present). E-F. *Avicularia rufa* Schiapelli & Gerschman, 1945 E. Hairiness (high score) in an adult female. F. Ontogenetic changes in color (present) in a juvenile. Arboreal habitat. Note the change in the abdominal pattern from the stage shown in E. Photos: Marlus Almeida (A), Caroline Fukushima (B, C, E), Ian Tarrant (D), and Flavio Terrassini (F).

**Table 1.**
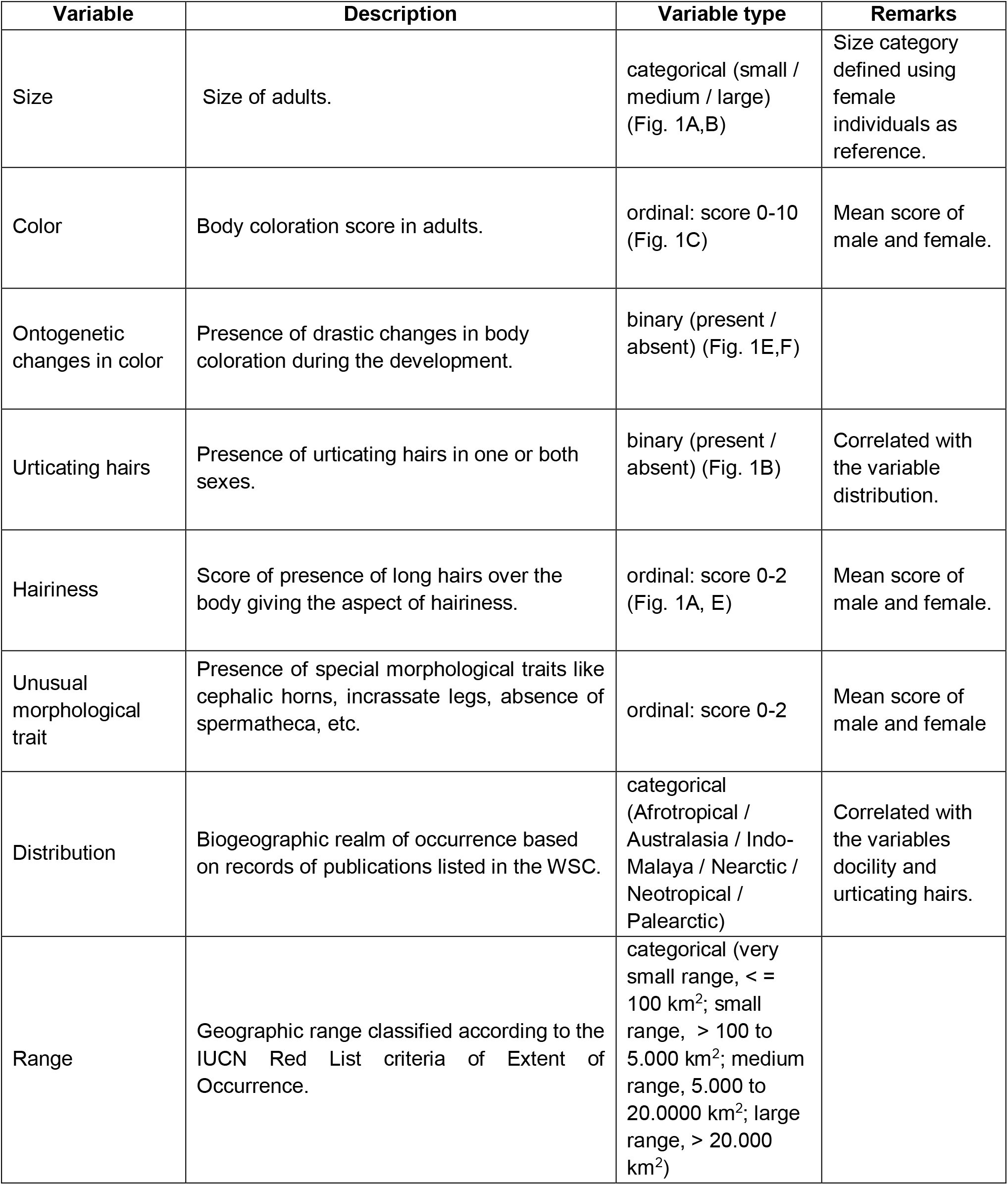
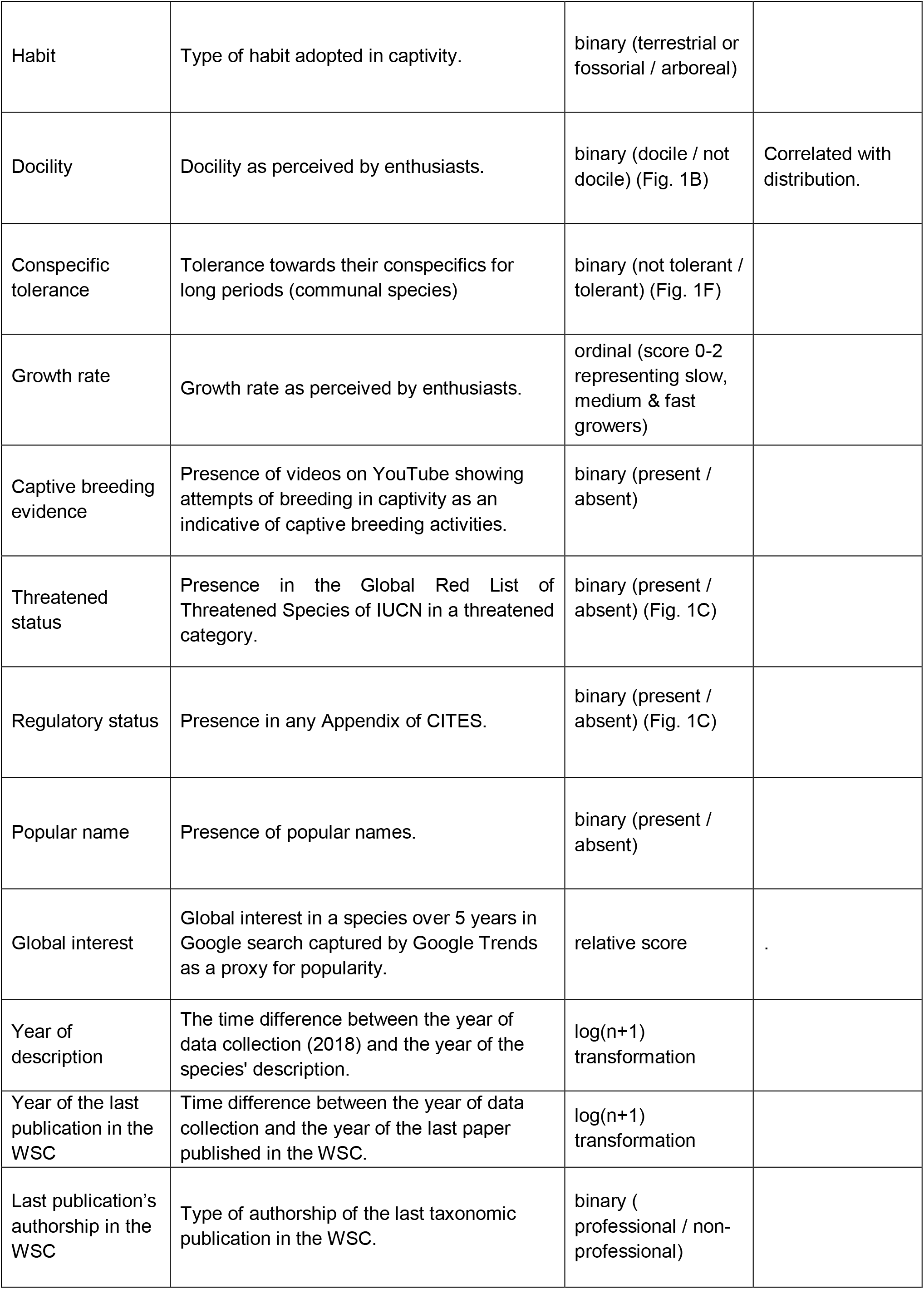
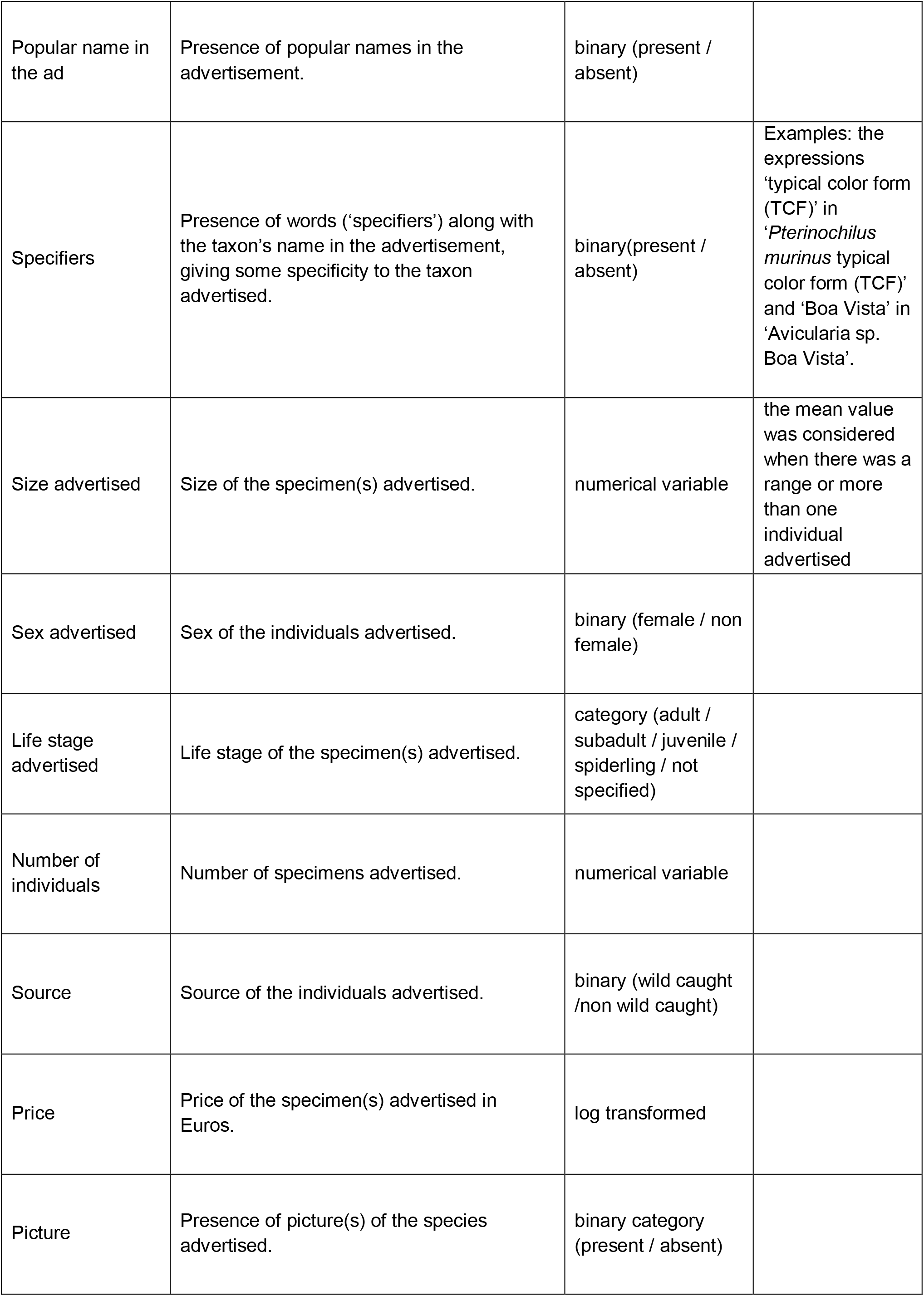
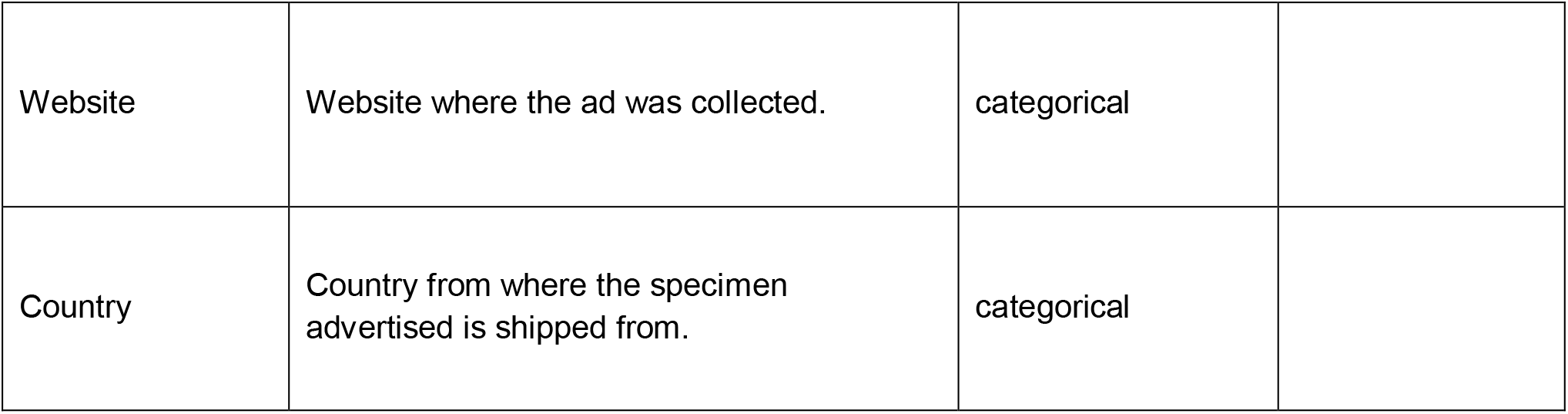
Species and advertisement attributes and correspondent description. For further details on trait description, units, data sources, and remarks see Appendix S1. CITES = Convention on International Trade in Endangered Species of Wild Fauna and Flora. IUCN = International Union for Conservation of Nature. WSC = World Spider Catalog v19.0.

| Variable | Description | Variable type | Remarks |
| --- | --- | --- | --- |
| Size | Size of adults. | categorical (small / medium / large) (Fig. 1A,B) | Size category defined using female individuals as reference. |
| Color | Body coloration score in adults. | ordinal: score 0-10 (Fig. 1C) | Mean score of male and female. |
| Ontogenetic changes in color | Presence of drastic changes in body coloration during the development. | binary (present / absent) (Fig. 1E,F) |  |
| Urticating hairs | Presence of urticating hairs in one or both sexes. | binary (present / absent) (Fig. 1B) | Correlated with the variable distribution. |
| Hairiness | Score of presence of long hairs over the body giving the aspect of hairiness. | ordinal: score 0-2 (Fig. 1A, E) | Mean score of male and female. |
| Unusual morphological trait | Presence of special morphological traits like cephalic horns, incrassate legs, absence of spermatheca, etc. | ordinal: score 0-2 | Mean score of male and female |
| Distribution | Biogeographic realm of occurrence based on records of publications listed in the WSC. | categorical (Afrotropical / Australasia / Indo-Malaya / Nearctic / Neotropical / Palearctic) | Correlated with the variables docility and urticating hairs. |
| Range | Geographic range classified according to the IUCN Red List criteria of Extent of Occurrence. | categorical (very small range, < = 100 km<sup>2</sup>; small range, > 100 to 5.000 km<sup>2</sup>; medium range, 5.000 to 20.0000 km<sup>2</sup>; large range, > 20.000 km<sup>2</sup>) |  |
| Habit | Type of habit adopted in captivity. | binary (terrestrial or fossorial / arboreal) |  |
| Docility | Docility as perceived by enthusiasts. | binary (docile / not docile) (Fig. 1B) | Correlated with distribution. |
| Conspecific tolerance | Tolerance towards their conspecifics for long periods (communal species) | binary (not tolerant / tolerant) (Fig. 1F) |  |
| Growth rate | Growth rate as perceived by enthusiasts. | ordinal (score 0-2 representing slow, medium & fast growers) |  |
| Captive breeding evidence | Presence of videos on YouTube showing attempts of breeding in captivity as an indicative of captive breeding activities. | binary (present / absent) |  |
| Threatened status | Presence in the Global Red List of Threatened Species of IUCN in a threatened category. | binary (present / absent) (Fig. 1C) |  |
| Regulatory status | Presence in any Appendix of CITES. | binary (present / absent) (Fig. 1C) |  |
| Popular name | Presence of popular names. | binary (present / absent) |  |
| Global interest | Global interest in a species over 5 years in Google search captured by Google Trends as a proxy for popularity. | relative score | . |
| Year of description | The time difference between the year of data collection (2018) and the year of the species' description. | log(n+1) transformation |  |
| Year of the last publication in the WSC | Time difference between the year of data collection and the year of the last paper published in the WSC. | log(n+1) transformation |  |
| Last publication's authorship in the WSC | Type of authorship of the last taxonomic publication in the WSC. | binary (professional / non-professional) |  |
| Popular name in the ad | Presence of popular names in the advertisement. | binary (present / absent) |  |
| Specifiers | Presence of words ('specifiers') along with the taxon's name in the advertisement, giving some specificity to the taxon advertised. | binary(present / absent) | Examples: the expressions 'typical color form (TCF)' in '*Pterinochilus murinus* typical color form (TCF)' and 'Boa Vista' in 'Avicularia sp. Boa Vista'. |
| Size advertised | Size of the specimen(s) advertised. | numerical variable | the mean value was considered when there was a range or more than one individual advertised |
| Sex advertised | Sex of the individuals advertised. | binary (female / non female) |  |
| Life stage advertised | Life stage of the specimen(s) advertised. | category (adult / subadult / juvenile / spiderling / not specified) |  |
| Number of individuals | Number of specimens advertised. | numerical variable |  |
| Source | Source of the individuals advertised. | binary (wild caught /non wild caught) |  |
| Price | Price of the specimen(s) advertised in Euros. | log transformed |  |
| Picture | Presence of picture(s) of the species advertised. | binary category (present / absent) |  |
| Website | Website where the ad was collected. | categorical |  |
| Country | Country from where the specimen advertised is shipped from. | categorical |  |

Given that some tarantula species exhibit sexual dimorphism, we calculated the mean value of a variable for both males and females in cases where the trait could vary (e.g., body color and hairiness). All numeric variables were standardized to have a mean of 0 and standard deviation of 1 during analysis, to ensure that large differences in the magnitude of different variable units did not affect coefficient estimates.

### Advertisement traits variables

We selected attributes that were present in most ads of the selected webstores (see Table 2 and Appendix S1 for mode details). The number of specimens available in stock was only displayed in three out of eight websites, therefore ads that did not display stock numbers were assumed to be selling single individuals.

**Table 2.** Frequency (%) of ad traits per website. Country codes for the web stores: CAN = Canada, GER = Germany, POL = Poland, SA = South Africa, UK = United Kingdom, USA = United States of America. N = 977 ads.

| Website | Ads | Pictures | Specifiers | Popular name | SEX |  |  |  | LIFESTAGE |  |  |  |  |
| --- | --- | --- | --- | --- | --- | --- | --- | --- | --- | --- | --- | --- | --- |
|  |  |  |  |  | Couple | Male | Female | Unsexed | Adult | Subadult | Juvenile | Spiderling | Unspecified |
| CAN | 20.6 | 19.4 | 5.3 | 0.8 | 0 | 0.4 | 1.1 | 19 | 0.2 | 0 | 4.6 | 15.8 | 0 |
| GER | 12.7 | 11.8 | 2.7 | 0.3 | 1 | 1.3 | 2.7 | 7.7 | 0.1 | 0.2 | 4.6 | 7.7 | 0.1 |
| POL1 | 25.4 | 20.1 | 7.4 | 13.1 | 3.6 | 1.7 | 5.7 | 14.3 | 2.6 | 0.6 | 14.2 | 7.9 | 0.1 |
| POL2 | 9.5 | 8.7 | 1.7 | 0 | 0 | 0.7 | 1.3 | 7.5 | 0 | 0 | 3.1 | 6.4 | 0 |
| SA | 7 | 6 | 0.5 | 6.9 | 0 | 0 | 2.4 | 4.6 | 1.7 | 0.1 | 3.3 | 1.8 | 0 |
| UK | 8 | 8 | 1.8 | 7.9 | 0 | 0 | 0.4 | 7.6 | 0.3 | 0.6 | 3.1 | 4 | 0 |
| **USA1** | 8.1 | 7.9 | 1.6 | 6.3 | 0 | 0.7 | 0.6 | 6.8 | 0.1 | 0 | 4.5 | 3.5 | 0 |
| **USA2** | 8.8 | 0 | 0.2 | 8.7 | 0.1 | 0.4 | 2.6 | 5.7 | 1 | 2.1 | 3.9 | 1.7 | 0 |
| **Total (%)** | 100 | 81.9 | 21.3 | 44 | 4.7 | 5.3 | 16.8 | 73.2 | 6 | 3.7 | 41.2 | 48.8 | 0.2 |

### Structural Equation Modeling

First, we used a mental model to express the hypothesized relationship among the variables (see Appendix S2 for further explanations). We used Structural Equation Models (SEMs) to study the network of relationships among species traits, advertisement traits, and price and abundance of OTUs (referred to as species hereafter) (Grace 2006). SEM is a powerful approach that can unite several statistical models for the analysis of datasets with multiple mutually intercorrelated dependent and independent variables (Lamb et al. 2011; Toomes et al. 2022). This approach models multivariate direct and indirect relationships by partitioning variable correlation between explanatory and response variables, allowing the representation of more complete hypotheses and revealing unanticipated relationships among variables (Schweiger et al. 2016, Toomes et al. 2022).

We adopted the piecewise SEM approach since the distribution of some of our variable relationships were non-Guassian (Lefcheck 2016), as determined by preliminary inspection of our data.

Some variables selected for this study only differed among species and not among ads, therefore abundance was not modeled at the scale of ads but instead summarized by species. We generated simple candidate gaussian and negative binomial generalized linear models (GLMs) to model price and abundance respectively, with each candidate containing one variable from the mental model. We chose generalized linear models with negative binomial distributions rather than Poisson distributions because preliminary investigation of candidate models revealed significant overdispersion. We included ads selling more than one individual of species when calculated abundance and average price (standardized to per individual). However, we omitted these ads from our ad-level analysis because some attributes such as sex could vary between individuals within the same ad.

### Construction of Initial SEMs

Preliminary investigation of linear regression models with price as a response variable showed that model assumptions (such as the normality of residuals) were not satisfied. As such, we natural log transformed prices and found assumptions to be satisfied (Appendix S3). We used Akaike’s Information Criterion (AIC) to select the best candidate model with the lowest AIC score (i.e., lower than the next-best candidate by an AIC difference > 5; see Burnham et al. 2011 for further details on AIC cutoffs). We started with univariate candidate models and added variables to the best candidate model in a stepwise fashion until there were no further improvements, as indicated by lower AIC score. This final model was used to construct an initial SEM path (see Appendix S4). In instances where multiple models of the same complexity were within ΔAIC > 5 of each other but still outperformed simpler models based on AIC, we continued adding variables to each leading candidate in a stepwise fashion until a single candidate model was selected. If there was not a single best candidate after three successive increases in model complexity, we selected the most complex model from which all best performing candidates stemmed from.

This approach resulted initially in two SEMs: one modeling price at the resolution of individual ads and one modeling both abundance and average price at the resolution of species. We consider that each country may have unique characteristics, policies, enforcement level, or other unobserved factors that can influence the observed data. By considering variable ‘country’, we can better capture and explain the complexities and nuances associated with the countries included in the analysis, which in turn can influence the patterns and variability in the price of ads. Similarly, each website may attract different subsets of customers and cater to different trading strategies (e.g., large-scale breeding of common species versus specialization in rarer species). Due to this, we performed an alternative version of the advertisement-level SEM that considered generalized linear mixed models (GLMMs) with the variables ‘country’ and ‘website’ as random effects, ultimately resulting in three SEM analyses.

For each SEM analysis, we constructed an initial path diagram based on the selected candidate models outlined previously. We also specified covariation between the following pairs of variables based on *a priori* knowledge of their interrelatedness (see remarks in Table 1 for details): presence of urticating hairs & docile behavior, presence of docile behavior & distribution category, distribution category and presence of urticating hairs. All candidate explanatory variables were considered in tests of directed separation (dSep tests hereafter), conditional on the existing specified relationships. We performed dSep tests on our initial paths to determine whether there was evidence to include additional relationships that improve model explanatory power. For further details of this process see Appendix S4.

The analyses were conducted in the R software version 3.4.4 (R Core Development Team 2019), with the PiecewiseSEM package to generate and evaluate SEMs (Lefcheck 2016). The explanatory variables were tested for collinearity prior to their final inclusion in the initial SEM path diagram using a variance inflation factor test in the car package (Fox & Weisberg 2018). Overdispersion was tested for using the AER package (Kleiber & Zeileis 2008). In case of collinearity, AIC values were compared and variables with the least explanatory power were excluded from the SEM (Shipley 2013). Relative variable importance scores for final models were calculated using dominance analyses with the domir package (Luchman 2023). See Appendix S3 for further details about the process.

## Results

We selected eight online stores across six different countries to collect the ads: the United States (n = 2), Canada (1), the United Kingdom (1), Poland (2), Germany (1), and South Africa (1). We obtained 977 ads after discarding duplicated or “out of stock” (4% of total ads), of which 855 were advertising only one individual animal.. A total of 261 scientific names (valid or not; 225 at species level) and 372 names (including scientific names in several taxonomic levels, popular names, variations, and different spellings/misspellings of a name) were present in the ads (see Appendix S5). We recorded 217 valid species and 83 “trade names” being advertised (see Appendix 6). Regarding source of the specimens advertised, 90.9% of the ads do not specify it, 4.7% are of wild-caught specimens, and 4.4 are of captive bred. For the frequency of other ad traits, see Tables 3–4.

**Table 3.** Number of species exhibiting binary trait variables. N= 225 species.= data not available.

| **Variable** | **Present** | **Absent** | **?** |
| --- | --- | --- | --- |
| Popular Name | 203 | 22 | 0 |
| Urticating Hairs | 123 | 102 | 0 |
| Ontogenetic changes in color | 38 | 187 | 0 |
| Habit (terrestrial) | 176 | 48 | 1 |
| Threatened status (IUCN-listed) | 6 | 219 | 0 |
| Regulatory status (CITES-listed) | 14 | 211 | 0 |
| Evidence of captive breeding | 171 | 54 | 0 |
| Last publication's authorship (non-professional) | 98 | 127 | 0 |
| docility (docile) | 87 | 125 | 13 |
| Conspecific tolerance | 21 | 202 | 2 |

**Table 4.** Number of species exhibiting the non-binary trait variables. N = 225 species.= data not available.

| Variable | Category | Number of Taxa | Variable | Score | Number of Taxa |
| --- | --- | --- | --- | --- | --- |
| Distribution | 0 | 26 | Color | 0 | 9 |
|  | 1 | 5 |  | 0.5 | 11 |
|  | 2 | 48 |  | 1 | 13 |
|  | 3 | 15 |  | 1.5 | 9 |
|  | 4 | 129 |  | 2 | 24 |
|  | 5 | 2 |  | 2.5 | 22 |
| Range | 0 | 8 |  | 3 | 29 |
|  | 1 | 36 |  | 3.5 | 15 |
|  | 2 | 70 |  | 4 | 17 |
|  | 3 | 109 |  | 4.5 | 10 |
|  | ? | 2 |  | 5 | 17 |
| Hairiness | 0 | 167 |  | 5.5 | 8 |
|  | 0.5 | 10 |  | 6 | 17 |
|  | 1 | 20 |  | 6.5 | 4 |
|  | 1.5 | 8 |  | 7 | 13 |
|  | 2 | 20 |  | 7.5 | 2 |
| Size | 1 | 33 |  | 8 | 3 |
|  | 2 | 157 |  | 8.5 | 0 |
|  | 3 | 35 |  | 9 | 1 |
| Growth rate | 0 | 49 |  | 9.5 | 0 |
|  | 1 | 49 |  | 10 | 1 |
|  | 2 | 107 | Unusual morphological trait | 0 | 218 |
|  | ? | 20 |  | 0.5 | 2 |
|  |  |  |  | 1 | 5 |

### Species Trait-level Model

A final SEM was selected (Fig. 2) with non-significant Fischer’s C value of 63.78 (df = 60, p = 0.345), indicating that no paths were missing that would improve SEM explanatory power. See marginal means plots for all categorical variables included in the final SEMs in Appendix S7, Fig. S7.1. No direct variable paths were removed from the initial SEM path diagram (see Appendix S4) during our analysis, and dSep tests supported the addition of direct effects of urticating hairs, breeding evidence and conspecific tolerance on average price. We found no evidence for covariation between price and abundance at the species level after accounting for the explanatory variables that directly influence price and/or abundance.

**Figure 2.**
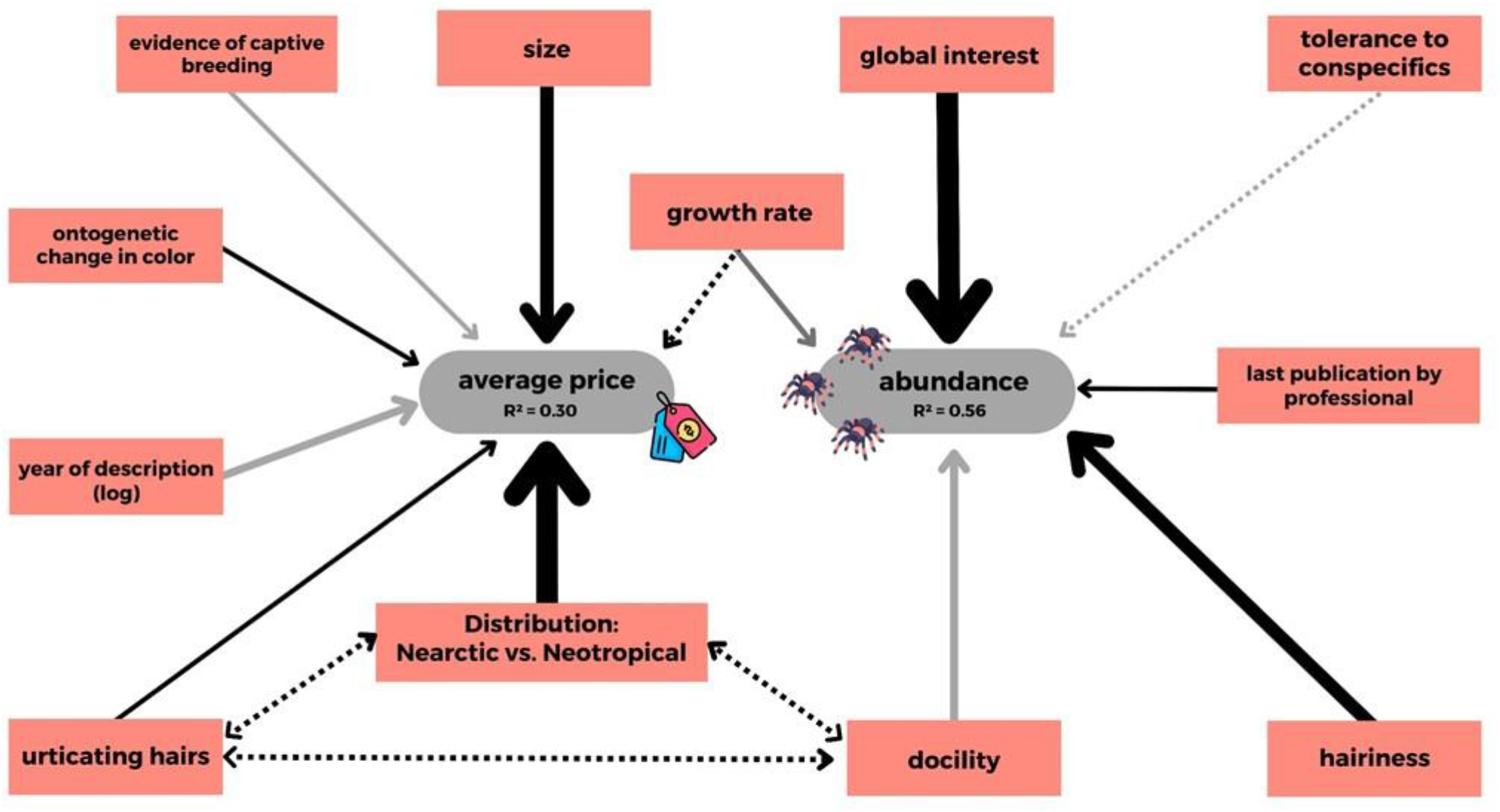
Final Structural Equation Model (SEM) path illustrating the relationships between tarantula species traits (orange boxes), price, and abundance in the online pet trade. The relative variable importance (RVI) scores are visually represented through arrow thickness, where greater thickness indicates higher importance. Positive effects are depicted by black arrows, while negative effects are indicated by gray arrows. Dashed arrows symbolize paths with non-significant effect size (p < 0.05), whereas non-dashed arrows represent paths with a significant effect size (p > 0.05). Covariation between variables is donated by double-ended arrows.

A high proportion of the variability in abundance was explained by the negative binomial GLM component of the final SEM (R^2^ = 0.56). Species were significantly more abundant in trade if they had higher global interest (0.868 ± 0.0735, P-value < 0.0001, see Fig. 3), slower growth (−0.181 ± 0.0767, P-value = 0.0186), are hairy (0.238 ± 0.717, P-value = 0.0009) and perceived as non docile (1.81 ± 0.122, P-value = 0.0008). Species with last publications authored by professional arachnologists (1.73 ± 0.120) were also significantly more abundant in the trade (P-value = 0.0086) than those published by non-professionals (1.35 ± 0.139). Global interest (69.9%) and hairiness (16.9%) were the most important variables driving variability in abundance. See Appendix S8 and S9 for specified relationships.

**Figure 3:**
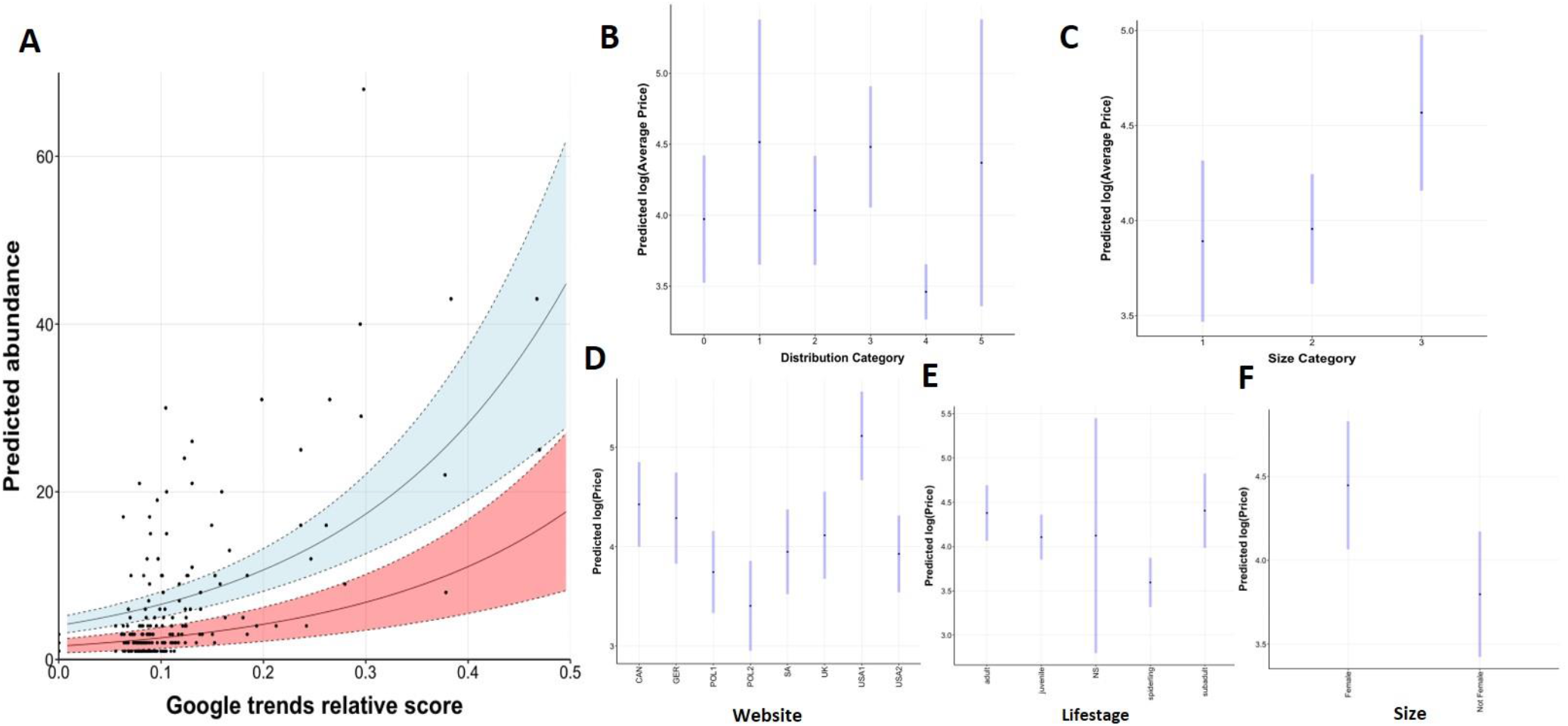
Model predictions showing relationships between (A) species trade abundance and global interest expressed by google trends relative scores. Raw data displayed as points. Predictions displayed in red (docile species with conspecific tolerance and last publications authored by non-professional arachnologists) and blue (non-docile species without conspecific tolerance and last publications authored by professional arachnologists) with hairiness score held at its mean value. Five data points with Google trends above 0.5 are omitted to aid viewability. (B) natural log(average price) and distribution category; (C) log(average price) and size category; (D) log(price) and website of advertisement; (E) log(price) and lifestage of advertised animal; (F) log(price) and sex of advertised animal. Predictions are marginal means of each category with all other variables held at their average value. 95% confidence intervals displayed in blue.

A reasonable proportion of the variability in average price was explained by the linear regression component of the final SEM (R^2^ = 0.30). Distribution category (43.6%) and size category (22.6%) were the most important variables driving variability in price (Appendix S9). See model predictions for these variables in Fig. 3. Species were significantly higher in average price if they were more recently described (i.e., time since description was lower; (−0.114 ± 0.0467, P-value = 0.0157, relative variable importance score = 11.2%), had urticating hairs (4.352 ± 0.208, P-value = 0.0046) and undergo drastic ontogenetic shifts in coloration (4.35 ± 0.215, P-value = 0.0046). Species in the large size class (4.57 ± 0.208) are significantly higher in price than ‘median’ (3.96 ± 0.215, P-value = 0.0013) and small species (3.89 ± 0.215, P-value = < 0.0001). Species that have captive breeding evidence were significantly lower in average price (−0.395 ± 0.137, P-value = 0.0046). Significant differences were detected between distribution categories (P-value < 0.0001), with Tukey’s pairwise comparisons revealing significantly higher prices (P-value < 0.0001) for Nearctic species (4.48 ± 0.217) than for Neotropical species (3.46 ± 0.0987) (see Appendix S10).

### Advertisement Trait Fixed-Effect Model

Our fixed-effects SEM analysis yielded a final path with non-significant Fischer’s C value of 13.75 (df = 22, p = 0.91), indicating that no paths were missing that would improve SEM explanatory power. The marginal means plots for all categorical variables included in the final SEMs can be seen in the Appendix S7, Fig.S7.2-3. The direct relationship between the size of advertised animals and price was removed during dSep analysis, whereas relationships with 13 variables were added, totalling 18 variables with direct relationships to price (see Appendix S4 for further details in specified relationship). The linear regression component of the final SEM explained a substantial portion of the variability in price (R^2^ = 0.59), more so than the equivalent model at the species level. Most variables had low relative importance (<6% each; see Appendix S9), with the highest scores for website (35.9%), lifestage (13.6%), sex (female/non-female; 11.4%) and source category (8.7%). See model predictions for these variables in Fig. 4. We found that ads of less abundant species are higher in price (−0.0104 ± 0.024), a direct relationship not detected at the species level. Additionally, ads had significantly higher prices if they featured species with low scores of hairiness (−0.0907 ± 0.0312, P-value = 0.0037), high color scores (0.093 ± 0.0319, P-value = 0.0036), and were newly described (−0.105 ± 0.0271, P-value = 0.0001). Ads displaying species with slower growth rates were slightly higher in price although the trend was not significant (P-value = 0.682). Price was higher for ads selling species with ontogenetic shifts, a trend that approached significance (P-value = 0.056).

Advertisements of species with smaller geographic ranges (log(price) = 4.21 ± 0.196) had significantly higher prices (P-value = 0.0265) than those displaying species with larger geographic ranges (4.00 ± 0.187). Ads of docile species (4.23 ± 0.189) had significantly higher prices (P-value = 0.0012) than ads of species considered more aggressive (4.02 ± 0.189). Significant differences were detected in the price of ads of species belonging to different size classes (P-value < 0.0001), with ads of large (‘giant’) species presenting the highest prices (4.44 ± 0.197) and ads of small (‘dwarf’) species presenting the lowest (3.76 ± 0.202). Similarly, prices varied in ads of different life stages (P-value < 0.0001), with ads of sub adults (4.41 ± 0.214) or adults (4.38 ± 0.160) higher in price than spiderlings (3.60 ± 0.142); and in ads of different sexes (P-value < 0.0001), with ads of female tarantulas (4.45 ± 0.194) having higher prices than non-females (unsexed or males; 3.80 ± 0.191). The source of the specimens advertised was also selected as a significant predictor in the model (P-value < 0.0001), with ads of captive bred individuals with the highest prices (4.76 ± 0.266). Ads had significantly lower prices if they displayed species: listed in CITES (P-value = 0.0033), with tolerance to conspecifics (P-value < 0.0001), with evidence of captive breeding (P-value < 0.0001), displayed a photo of the species being sold (P-value = 0.0214), or used a popular name to identify the animal (P-value = 0.0015). Finally, significant differences on the price in the ads were detected between websites (P-value < 0.0001), with ads from an e-commerce from Poland (POL2, 3.41 ± 0.230) displaying the lowest prices and ads from an e-commerce from the USA (USA1, 4.12 ± 0.223) with the highest prices. Pairwise comparison tests revealed that advertisements on USA1 were significantly higher than all other sites, including from the same country (Appendix S10). Similar to the previous SEM findings, our observations indicated that the distribution category and the presence of urticating hairs potentially exerted an indirect influence on advertisement price, mediated by their covariation with “docility.”

### Advertisement Trait Mixed-Effect Model

Our alternative advertisement-level SEM using linear mixed effect modeling yielded a final path with non-significant Fischer’s C of 19.51 (df = 22, p = 0.614), with a high proportion of the variability in price explained by the model (R^2^ = 0.57). As the advertisement-level SEM with no random effects presented smaller Fischer’s C values and higher R-squared values, it was selected as the preferred model, though the mixed-effect SEM may supplement our insights into price-abundance-trade relationships.

Consideration of candidate mixed models supported the inclusion of ‘country of export’ as a random effect (see Appendix S8 for further details in specified relationship). The magnitude, direction and statistical significance of most variables were similar between the fixed effect and mixed effect final SEMs (see Appendix S8 for specific model output) but the inclusion of random effects led to the loss of a direct relationship with range and source categories and the inclusion of three additional variables with direct effect on advertised price. For two of them the trends were not significant, namely urticating hairs (P-value = 0.0611), with ads of species with urticating hairs having higher prices, and size of the specimen advertised (P-value = 0.4897), with a negative effect in the price of the ads. However, significant differences were detected between distribution categories (P-value = 0.002), with ads with price highest for Australasian and Nearctic species and lowest for Neotropical species, as observed in the species-level SEM. Our dSep tests provided evidence for the addition of websites of advertisement and therefore we considered this variable as random effect. However, inclusion of websites as random effects led to singularity issues (likely resulting from only small differences in response occurring between most sites) so we included the variable as a fixed effect instead.

## Discussion

Our results support the hypothesis that species traits are prominent drivers of price and abundance of tarantulas in the pet trade. However, we found no evidence for price-rarity dynamics at the species level unlike previous research in other taxa (Su et al. 2015). This implies that the availability of species in trade and their market value are driven by independent processes, as growth rate was the only variable in our models with direct relationships to both responses. In a comparable manner, Shivambu et al. (2020) found that price is not a good predictor for species availability in the South African tarantula pet trade. At the resolution of individual advertisements, there was evidence for a small effect of species abundance on price. As typical in competitive market dynamics, increasing offers drives prices down. However, this variable has low relative importance (<5% of the total contribution to model explanatory power) and does not strongly support the presence of price-market rarity dynamics.

### The role of morphological, ecological, and behavioral traits

Tarantulas are the world’s largest spiders (Montes de Oca & Mendoza 2020) and thus it was expected that their size would be an attractive trait in the pet trade. The preference for larger body size species is a common trend in pet trade: for instance, it is present in the mantis (Battiston et al. 2022), reptile (Toomes et al. 2022), and amphibian trade (Mohanty & Measey 2019).

Tarantulas with long hairs covering their bodies possess an aesthetic appeal that may increase empathy towards them. They may be perceived as “cute”, increasing the desirability of these species as pets, as has been expressed by consumers of wildlife-related social media content for charismatic taxa (Moloney et al. 2021). Alternatively the hairy appearance of some tarantulas could enhance their resemblance to vertebrates and this would increase the affinity towards a species, as this may be heavily influenced by the degree of similarity to humans (Batt 2009).

Color-related terms are often used in the common names of tarantula species (Marshall et al., 2022), highlighting the significance of this trait in determining their overall attractiveness. Similar trends can be observed in other trade groups, such as birds (see Senior et al., 2022). Color morphs, in certain cases, could even represent potentially undescribed species (Marshall et al. 2022). Despite never being tested before, body coloration is considered an important trait on the purchase of tarantula species (Marshall et al. 2022), though our analysis suggests its importance is relatively low.

Certain behavioral traits have shown significance in determining species abundance in the market. Ads of docile species are higher in price, and this may be due to the fact that docile species are preferred by those beginning in the hobby (Montes de Oca & Mendoza 2020) and/or may be easier to handle and care for. On the other hand, species perceived as not docile or that do not possess high tolerance for conspecifics tend to be more abundant in the market. The higher abundance of more aggressive and less conspecific tolerant species may be connected to people’s perception of spiders as intimidating and dangerous creatures evoking fear and apprehension, but also some fascination (Mendoza 2020), a feeling that could be a motivational driver for enthusiasts. Lassaline et al. (2023) found that the majority of the most popular traded invertebrates in Australia are dangerous to humans, suggesting a disproportionate desire for challenging exotic pets.

### Extrinsic attributes

The popularity of a species, as indicated by the score of global interest obtained from Google Trends data, is the main driver of market abundance. Trend scores are not driven by trade stakeholders alone but also by the general public, including prospective pet owners. It is unclear whether popular species online are targeted by traders or if species available in the market become popular with the public, but probably both factors act in a positive feedback loop to drive the market. Future research could investigate temporal trends in online popularity and trade abundance to determine whether predictable time-lags exist between a rise in species popularity and its widespread trade. If popularity drives trade, as assumed in our model, Google Trends may provide a valuable early warning indicator of the emergence of new species in online markets, facilitating a more proactive approach to tarantula trade management. It should be noted, however, that our proxy for popularity will not be generalisable to all jurisdictions since Google is not the only major search engine in some areas (e.g., China) (de Oliveira Caetano et al. 2023) and that analogous measures of online popularity should be considered.

There is a clear incentive to trade newly described taxa as previously suggested by Marshall et al. (2022), a trend also found for reptiles and amphibians (Altherr and Lameter 2020). This novelty-price relationship is evident in a recent controversial case where breeders collected and potentially smuggled a new tarantula species from Malaysia, leading to its scientific description; notably, this newly discovered species was being sold for high prices (US$300) through their online pet shops based in Poland and the UK (Law 2019). Our findings suggest that the desire for novelty is generalisable to any newly described tarantula. Such species are unlikely to have been assessed for their conservation status and the extent of their distribution may not be well documented. As such, it is extremely difficult to determine the potential effects of wild harvesting on their populations, and it is concerning that their market value is higher during the period of time where they are most poorly understood.

Although large quantities of traded tarantulas come from the wild (Marshall et al. 2022), ads selling captive bred spiders showed significantly higher prices than ads selling wild caught species, different from the findings in turtle trade in Hong Kong (Sung & Fong 2018), but similar to the general results for exotic pets found by Hausmann et al. (2023). Higher prices can be linked to higher costs to breed and keep animals in captivity to be posteriorly sold. As such, the willingness of prospective pet keepers to a more sustainably sourced (i.e., captively bred) pet may be influenced by the increased cost. In such cases, establishing captive breeding populations may not eliminate the demand for wild-caught pets. Therefore, before implementing this conservation tool, careful technical, legal, and ethical considerations are necessary, especially those pertaining to sustainability, demand, and legality of captive bred specimens derived from illegally collected parents (see CEC 2017 and Fukushima et al., 2021). And if adopted, captive breeding regulations must ensure equitable benefits for the species’ native countries and be accompanied by awareness campaigns to promote informed consumer choice.

Our findings in the lack of any direct effects of CITES or IUCN status on price or abundance at the species level has important implications for the future management and regulation of the international tarantula trade. For example, if trade in a given species decreases (e.g., is banned or reduced for conservation or biosecurity purposes), the perceived value of that species may not substantially differ, as observed in the global pet trade in amphibians (Mohanty & Measey 2019) and raptors (Nijman et al. 2022). The ongoing trade suspension of all Mexican CITES-listed species (which includes the popular *Brachypelma* and *Tliltocatl* species) since March 2023 provides a unique opportunity to observe and analyze the tarantula market’s response to global bans.

### Individual attributes

The choice of which pet to purchase is sometimes dependent on fine-scale attributes of the individual animal. Unsurprisingly we found direct effects on price from several variables that differ animal-to-animal. The life stage of the advertised tarantula specimen emerges as the second prominent factor contributing to the variability in prices among ads. Spiderlings are often priced lower due to their minimal time and resource investment before sale, unlike adults or subadults, which may require years to reach that stage. In the case of wild-caught specimens, higher prices may result from the challenges of locating, collecting, and transporting (legally or illegally) species in later life stages.

Ads displaying females had significantly higher prices, as expected. Females may be more attractive because they tend to have larger body sizes, and longer lifespans than males (Montes de Oca & Mendoza 2020). Captive records indicate that females can attain lifespans of 20-34 years (Montes de Oca & Mendoza 2020), which surpasses the life expectancy of more common pets such as dogs (Teng et al. 2022).

### External factors to consider

While there are intrinsic species and individual traits that influence desirability in trade, a variety of external contextual factors are also important to consider. Chiefly, variations in the average price of tarantulas are more associated with the specific website of sale than to variables such as source, market abundance, presence in CITES or species specific traits. Notably, our findings reveal that the two e-commerce platforms displaying ads with the highest prices are located in North America. Differences in consumer base and expendable income as well as in regulatory framework and in the level of compliance may explain this difference. For instance, the USA has implemented stringent laws and policies (chiefly the Lacey Act, that prohibits trade in species that are protected by the domestic legislation in their country of origin) to protect both domestic and foreign species. As such, prices may be higher due to the costs involved in collecting, transporting, importing species, and in adhering to specific regulations. On the other hand, the two e-commerce platforms with ads at the lowest price are situated in Poland. The country has gained a reputation in the tarantula hobby community for its collectors and breeders who travel abroad to acquire and breed specimens to feed the international pet market (e.g. in Law 2019; Fukushima pers. obs.). The fact they may be one of the suppliers in the beginning of the trade chain may explain the low prices in the ads. The low prices along with the fact that one of the Polish shops display prices also in Euro and US dollars (the national currency being the zloty) suggest the idea that they largely rely on international transactions.

## Challenges and Recommendations

This study offers a snapshot of the tarantula trade, despite constraints due to language barriers and restricted market access in regions like the Middle East and Asia. Additionally, as it represents only a portion of overall trade, there may exist discrepancies in species availability and prices compared to physical stores (Shivambu et al. 2020). Thus, careful consideration is needed when interpreting and generalizing the findings, especially when translating them to broad conservation actions.

A considerable proportion of trade takes place using ‘trade names’. Evidence suggests that tarantula identification accuracy in the market is inadequate (Shivambu et al., 2020). These factors contribute to enhancing the already large taxonomic confusion of the group (see Fukushima et al. 2020), also hampering the study, regulation, monitoring, and management of its trade.

No websites in our study provide any evidence of the provenance or legality of the products they sell, except for one store that stated to provide CITES permit numbers for some of the CITES-listed species advertised. The vast majority of the ads (90.1%) does not display the source of the individual(s) advertised. The current state of underregulation in both the invertebrate pet trade and e-commerce presents potential scenarios where online pet stores can thrive on illegal and unsustainable practices. This concern is exemplified by the arrest of an online pet store owner attempting to smuggle around 1,000 tarantulas from South America to the UK for sale in his establishment, where many died due to poor transport conditions (Panther 2015). Such instances highlight the need to address animal welfare, sustainability, transparency, and traceability in the tarantula trade (CEC 2017).

We suggest stronger efforts in regulating and monitoring the trade in countries that may act as key importers and exporters of tarantulas worldwide such as Poland. For instance, priority can be given to the adoption of the new Digital Services Act in the EU, that will equip Member States with tools to tackle the escalated illegal online trade and the boom of e-commerce (Paquel 2016). Additionally, it could adopt a national positive list on wild and exotic animals to be kept as pets, as suggested by the revision of the EU Action Plan Against Wildlife Trafficking (European Commission 2022). However, while there is evidence supporting the efficacy of positive lists in regulating the pet trade (Toland et al. 2020), it is important to consider the potential backlash from exotic pet enthusiasts, as it may result in an increase of engagement in illegal trade. As a significant consumer region in the pet trade, Europe bears the responsibility for conserving species beyond its geographical boundaries (Auliya et al. 2016). On a global scale, urgent action is required to regulate exotic pet e-commerce platforms, emphasizing the importance of transparency in providing accurate information regarding the origins, sustainability, and legality of the wild animals available on their websites.

### Stakeholders and tools for the way forward

Hobbyists and their societies play a role in enhancing the popularity of specific tarantula species and in disseminating species discoveries and scientific papers in taxonomy. Importantly, our research demonstrates that these factors have a noteworthy influence on the variation in market price and abundance of tarantula species. Thus, they can influence consumer demand and trends in the market (Jain et al. 2021). Furthermore, tarantula societies have the potential role in providing information and raising awareness among their members about the legal implications of the trade. Yet in some settings, the ad-hoc keeping of animals by independent collectors is sometimes perceived to be part of species conservation even when there is no coordination among breeders or between them and organizations working in the natural habitat of those species, making it unlikely that any positive conservation benefits are materialized. It is thus key that conservationists more fully engage with hobbyist societies in order to change prevailing social norms around the relationship between pet keeping and species conservation. Indeed, the potential for these organizations to serve as a hub for consumers in the tarantula trade makes them “institutional influencers”, which could be leveraged to effectively deliver messages to a large proportion of consumers (Olmedo et al. 2020). Given the history these organizations have within hobbyits communities they may well be the most strategic messenger for conservation messaging trying to reach consumers in the tarantula pet trade. Consumer research with consumer groups is a high priority to continue to explore how they can most effectively be reached and influenced (Veríssimo et al. 2020).

The use of data from online ads remains an overlooked and underutilized resource for understanding the dynamics of supply and demand in the wildlife trade. Online ads can serve as a valuable tool for real time monitoring and understanding the exotic pet trade as they reflect the understanding of suppliers, for example regarding the information that buyers find relevant. This knowledge is central to maintaining a successful business and so is grounded and tested by the sellers own personal experience, with only those that provide relevant information to buyers staying in business in the long term. The information in ads can also be instrumental in the design of consumer-facing interventions (e.g. o help design the messaging for social marketing campaigns to reduce consumer demand), having the potential to inform the design and framing of messages with a higher likelihood of being relevant to consumers and therefore of being noticed and eventually effective at influencing consumer behavior. By leveraging the marketing insights of those with the most knowledge of consumers in the pet trade conservationists can improve their own chances of effectively influencing consumers, a key step if the trade is to move away from threatened species and specimens collected in unregulated settings and towards sustainability, through responsible ownership and a regulated trade with clear benefit sharing mechanisms that can bring benefits to those living alongside the wildlife.

## Acknowledgements

We would like to thank Ernie Cooper, Jorge Mendoza, Martin Gamache, Chris Hamilton, Rodrigo Orozco, Antonio Tosto, and numerous others for generously sharing their knowledge about tarantulas; and Caio Roza for suggestions on the manuscript. Our special gratitude to Rick West, a dedicated advocate for tarantula conservation, whose patient guidance about the subject has been invaluable throughout the years. We thank Flavio Terrassini, Marlus Almeida and Ian Tarrant for providing great pictures. We thank the Oxford Martin School for granting a fellowship for visiting scholars at the University of Oxford in the UK facilitating valuable academic exchanges among the authors. This research was supported by the Kone Foundation.

## Conflicts of Interest

The authors declare no conflicts of interest.

## Supporting Information

Supporting Information for Traits (Appendix S1), Mental Model (Appendix S2), Q-Q plots (Appendix S3), SEM analysis: dSep Tests and Initial Paths (Appendix S4), List of Taxa in the ads (Appendix S5), Taxa and ‘trade names’ advertised per website (Appendix S6), Marginal Means Plots for Categorical Variables (Appendix S7), SEM analyses: Final Paths (Appendix S8), Relative Variable Importance Scores (Appendix S9), and Tukey’s pairwise comparison tests (Appendix S10) are available online. The authors are solely responsible for the content and functionality of these materials. Queries (other than absence of the material) should be directed to the corresponding author.

## Notes

### Competing Interest Statement

The authors have declared no competing interest.

