## Supplementary Material for "A big and hairy business: Investigating the interplay of species traits and trade dynamics in the tarantula pet market"

**APPENDIX S1 -** *Traits*

Taxonomy is based on the World Spider Catalog v.19.0 (herein WSC). We consider the information available until August 2018.

1. **Species Traits**

**Size**

Size in adults of a species, based on size of female individuals.

While it is generally true that females are larger than males in most tarantula species, both sexes tend to reach sizes of similar categories as adults (Hénaut et al. 2015). We consider size a categorical variable, classified as small ('dwarf'), medium, or large ('giant'). Large tarantula species are called 'giant’ species in tarantula hobby, as they can achieve substantial sizes like *Thepahosa stirmi* Rudloff and Weinmann 2010, which females can reach a body length up to 10 cm and leg span of 27cm, and males body length up to 9 cm and 26-28 cm of leg span (Montes de Oca & Mendoza 2020). The smallest theraphosid spiders are known in the hobby as ‘dwarf’ species, such as species of the genus *Cyriocosmus* Simon 1903, which can present body lengths around 3 cm and leg span around 5 cm for females, and 1.5 cm of body length and leg spans smaller than 5 cm for males (Fukushima et al. 2005). The classification adopted in the variable is based on how hobbyists categorize the species, with some having 'dwarf' or ’pygmy’, or 'giant' in their popular name. Information regarding that or the average size of adults was taken primarily from accessory texts published in the ads, or information in the species description. If the information was lacking in these sources, we searched in other web markets, and in breeders’ or enthusiasts’ blogs, or other online sources (e.g., Arachnoboards, The Tarantula Collective). Although the information sourced from enthusiasts may lack scientific validation and is primarily based on experiential knowledge, it remains relevant for our study objectives as it represents the size of adult specimens as perceived by the tarantula enthusiasts. Species with females reaching more than 17 cm of leg span were considered large species; species with females reaching 5-6 cm of leg span were considered as small. Species with females reaching sizes in between were considered medium species. Quantitative measurement of size was not feasible due to various factors: female theraphosids can continue growing after reaching maturity (unlike males) (Montes de Oca & Mendoza 2020), size can vary based on factors such as diet and husbandry practices (Montes de Oca & Mendoza 2020), and the size mentioned in species descriptions often refers to juvenile or immature specimens (see Fukushima & Bertani 2017 for many species previously classified as *Avicularia* Linnaeus 1758). Additionally, there is limited reliable data available on tarantula body size since scientific papers typically include measurements only for the type species. Categorical variable: small, medium, or large.

**Color**

Score of body coloration of adults of a species.

To quantify the general coloration of a species, a scoring system from 0 to 10 was employed, being 10 the most colored tarantula species. The score was calculated by evaluating five factors with 0 to 2 points: carapace color, abdomen color, leg color, iridescence, and the presence of unusual contrasting colors (blue, red, orange, yellow, and purple). The sum of the points of each factor was considered the overall score of body coloration of the male or female of a given species. Since sexual dimorphism exists in certain tarantula species (Montes de Oca & Mendoza 2020), separate scores were assigned to males and females and the median value of these scores was then used as the overall score for the species. Although we have developed a score system to enhance reproducibility of results, we acknowledge a certain subjectivity in the evaluation of scores.

Ordinal variable, qualitative: score 0-10.

**Ontogenetic color change**

Presence of drastic ontogenetic changes in body coloration.

Many tarantula species can present changes in coloration during the development (Fukushima & Bertani 2017). Thus, spiderlings and juveniles can exhibit very different colors and patterns when compared to adults. Primarily information was sourced in scientific publication; in the absence of that, photographies of specimens in different life stages were examined to determine the presence or absence of drastic changes. Changes in coloration were considered drastic when colors (e.g. in *Theraphosa stirmi*) and carapace/abdominal patterns (e.g. in Aviculariinae species) differ among juveniles and adults. Binary variable: present or absent.

**Urticating hairs**

Presence of urticating hairs in one or both sexes of a species.

Urticating hairs are modified setae located on the abdomen or pedipalps of ‘New World’ theraphosid spiders (Kardeka 2019) and it is defense strategy against vertebrate and invertebrate predators and intruders (Cooke et al. 1972). When released, they can cause skin irritation and other allergy symptoms in humans (Castro et al. 1995), and can lead to various health issues (Bernardino & Rapuano 2000; Jalink & Wisse, 2021). The presence of urticating hairs is correlated with the distribution as only neotropical and nearctic species possess urticating setae (Bertani & Guadanucci 2003). It is generally assumed in the arachnoculture that species lacking these hairs tend to exhibit more aggressive displays towards humans (Montes de Oca & Mendoza 2020). Binary variable: present or absent.

**Hairiness**

The trait evaluated the 'hairiness' of a tarantula species, which refers to the presence of long hairs on their bodies.

Hairiness is considered a potential factor that increases the attractiveness of a species among buyers, as observed through discussions among enthusiasts on various forums. Male and female of species received a score from 0 to 2: 0 for spiders without very long hairs covering the body (e.g. in *Acanthoscurria geniculata*), 1 for spiders with some long hairs covering the body (e.g. in *Avicularia purpurea*), 2 for spiders with abundant long hairs in the body (e.g. in *Tliltocatl albopilosum*). In order to account for sexual dimorphism, the median value of hairiness scores for males and females was calculated and used as the overall score for the species. Although we have developed a score system to enhance reproducibility of results, we acknowledge a certain subjectivity in the evaluation of scores.

Ordinal variable: score 0-2.

**Unusual morphological trait**

Presence of uncommon morphological characteristics in a species.

The presence of uncommon structures like cephalic horns (e.g. in *Ceratogyrus* spp) or incrassate legs (e.g. in Citharognathus tongmianensis) was considered, as well as the lack structures that are present in the vast majority of spiders such as the spermatheca (e.g. in *Encyocratella olivacea*). Since sexual dimorphism exists in certain tarantula species (Montes de Oca & Mendoza 2020), separate scores were assigned to males and females and the median value of these scores was then used as the overall score for the species.

Ordinal variable: score 0-2.

**Distribution**

Biogeographic realm of the occurrence of a species, based on records of publications listed in the WSC.

This variable is related with urticating hairs and docility (see variables for further explanations). Distribution and venom potency may also be connected. ‘Old World’ species are considered as having stronger venoms than theraphosid spiders from Americas (Escoubas and Rash, 2004). Even though there is no record of human deaths resulting from tarantula bites, it is clear that some venoms are more toxic to humans than others and can cause persistent discomfort for several days (Garcia-Arredondo et al., 2015).

Categorical variable: Afrotropical, Australasia, Indo-Malaya, Neotropical, Palearctic.

**Range**

The geographic range of a species was primarily determined based on the literature available at the WSC, and was classified according to the IUCN Red List criteria of Extent of Occurrence (IUCN 2023). Extent of Occurrence (EOO) is defined as the area enclosed within the shortest continuous imaginary boundary that encompasses all known, inferred, or projected locations where a taxon is currently found, excluding cases of vagrancy (IUCN 2023). The choice of using this range measurement is primarily due to the limited availability of information regarding species occurrence, which is common for most species of tarantula spiders. The primary source for geographic occurrence of a species was the WSC. For species with limited information on their occurrence in the literature available at the WSC or discrepancies between their reported occurrence and actual localities, additional verification was conducted using INaturalist for checking for errors. Species known solely from their type locality were considered to have a very small range. The lack of information regarding species occurrence also increases the difficulty of obtaining them and subsequently raises prices. Information on the EOO was collected in the assessment of the IUCN Red List when possible; for the majority of species the area was calculated using Google Earth Pro v7.3.4. tools. Ordinal variable: very small range (< = 100 km2); small range (> 100 to 5.000 km2); medium range (5.000 to 20.0000 km2); large range (> 20.000 km2).

**Habit**

Type of habit in captivity.

Tarantulas can have terrestrial or arboreal habits and the type of habit adopted in captivity can affect the terrarium setup, some husbandry requirements, and care needed to keep a species as pet (Montes de Oca & Mendoza 2020). Information was primarily obtained in scientific papers. *Pachistopelma* species and their bromelicola habit (Bertani, 2012) were considered arboreal species since it is the primitive condition inside the Aviculariinae clade they are placed (Bertani, 2012). Binary variable: terrestrial or arboreal.

**Docility**

Docility of a species as perceived by enthusiasts.

Scientific studies specifically focusing on this trait are currently unavailable. However, the common presence in ads of information regarding docility may suggest its relevance in influencing the purchasing decision. Information regarding species' docility was explicitly stated or indirectly indicated through phrases such as "for beginners" or similar expressions. The term "aggressive" was considered the opposite of "docile" for the purposes of this study. In cases where docility information was not provided in the advertisements, we conducted additional searches in popular blogs, forums, other advertisements, or sought insights from breeders or taxonomic experts to gather relevant data. This comprehensive approach aimed to gather as much information as possible for the classification of species based on perceived docility since it ultimately drives the selection of a particular species. It is noteworthy that this observation can be linked to factors such as the species' distribution and the absence of urticating hairs. Enthusiasts often perceive "Old World" species (lacking urticating hairs) to be more aggressive (prone to bite) or skittish, which may influence their evaluation on this variable. Binary variable: docile or non docile.

**Conspecific tolerance**

Tolerance of a species towards their conspecifics.

Tarantulas are in general solitary species. However, some species can tolerate other conspecifics for long periods (some even being considered communal species, living peacefully with dozens of individuals). This characteristic may affect the attractiveness of a species. Additionally, in order to have specimens in communal set ups, enthusiasts need to buy several individuals. Information was collected in blogs, forums, in ads, or given by breeders or experts in the taxa. Species were considered with high tolerance when commonly reported among the hobbyist community as being able to live in communal set ups. Binary variable: tolerant or not tolerant.

**Growth rate**

Growth rate of a species as perceived by enthusiasts.

Given the scarcity of scientific data on species growth rates, we turned to webstore ads as potential sources of information, as they often provide small text descriptions of species traits found in these advertisements. Typically, growth rate information was not presented in terms of years required to reach maturity but rather in relative rates (slow, medium, fast). In instances where such information was missing, we conducted additional searches in blogs, forums, ads from other stores, or consulted breeders or taxonomic experts to obtain growth rate data. We classified species based on how they are perceived by buyers, as it is ultimately this perception that shapes their purchasing choices. In a conservative approach, we adopted the slowest rate when growth rates were described in between two categories. For example, if a species was described as having a medium-fast growth rate, we classified it as medium-paced. This approach minimized potential bias and allowed for a more objective categorization of species based on available information. Ordinal variable: slow, medium, or fast.

**Captive breeding evidence**

Presence of evidence of captive breeding activities of a species given by presence of videos on YouTube.

Reliable documentation on captive breeding in tarantulas is scarce. Thus evidence of captive breeding was collected on videos on YouTube. The platform is the largest global online video website and it is considered to be one of the most influential social network platforms, playing also a role on the exotic pet trade (Moloney et al. 2021). The search methodology involved querying YouTube using the search string 'mating' AND '(scientific name of species)' (e.g., mating AND 'Acanthoscurria chacoana'). We analyzed the content of the first 30 videos posted until July 2018, and we considered as evidence of captive breeding activities any video featuring mating attempts, regardless of success. To minimize distractions and focus solely on relevant content, the browser extension "unhook" was utilized. This extension concealed YouTube-related videos, comments, video suggestions, homepage recommendations, and the trending tab. Consequently, the number of unrelated videos appearing in the search was reduced, and the identification of mating videos featuring the target species was enhanced. Binary variable: present or absent.

**Threatened status**

Presence of a species in the Global Red List of Threatened Species of IUCN (the International Union for Conservation of Nature) in a threatened category (critically endangered, endangered or vulnerable) by June 2018.

Presence and status of tarantula species were searched in the IUCN website https://www.iucnredlist.org/

Binary variable: present or absent.

**Regulatory status**

Presence of a species in any Appendix of CITES (the Convention on International Trade in Endangered Species of Wild Fauna and Flora) by June 2018.

Status of the species were checked using the tool Species+ (https://speciesplus.net/).

Binary variable: present or absent.

**Presence of a popular name**

Presence of a popular name in English for a species within the hobby.

We limited our analysis to popular names in English due to the predominant use of this language in international forums and communities of tarantula enthusiasts, which is likely to capture global usage rather than regional variations. Initially, the collected advertisements were examined to determine whether a popular name was mentioned. If no popular name was found in the collected ads, we conducted additional searches using the Google search engine and the Boolean search string ‘(species' scientific name)’ AND ‘popular name’. To establish a species as having an established popular name in the hobby, we required the presence of the English popular name in at least three different sources obtained from the first three pages of the search results. Species that lacked an English popular name in the hobby typically yielded search results of fewer than three pages on Google, enabling a comprehensive examination of all content to ensure the absence of commonly used popular names. Binary variable: present or absent.

**Global interest**

The global interest in a species over a 5-year period was measured using Google search data obtained from Google Trends, and this interest was compared to that of the species *Poecilotheria metallica*.

This metric can be seen as a proxy for species popularity. Google Trends is a platform that provides data on the relative volume of Google searches for a specific term within a defined time period and region of interest (Correia 2018). To ensure fair comparisons between different terms, Google Trends normalizes the search data using a scale of 0 to 100 (refer to https://support.google.com/trends/answer/4365533?hl=en). First step was to select a comparison term. Prior analysis of Google Trends data indicated that *Poecilotheria metallica* was the species with the highest popularity peak during the specified time period. Hence, we selected it as the reference term for comparing the global interest in other species (e.g., ‘Poecilotheria metallica’ versus ‘Avicularia avicularia’). The Google Trends settings were configured to conduct global searches from 1 August 2013 to 23 July 2018, spanning a total of 260 weeks. The terms used in the searches were validated following the methodology outlined by Correia (2018). The search results were exported to a CSV file, and the search volumes of the species were summed. In our study, the variable value represents the worldwide search volume of a given species relative to the search volume for the term 'Poecilotheria metallica'. The calculation for the global interest score is as follows: Global interest = Sum of search volume of a species / Sum of search volume of *Poecilotheria metallica*. The resulting score represents the variable value used in the analysis. Ordinal variable.

**Year of Description**

The lag (Δ) represents the time difference between the year of data collection (2018) and the year of the species' description.

We consider the year of description as informed on the World Spider Catalog v.19.0. When dealing with subjective synonyms (i.e., names based on different holotypes), we considered the year of description for the name used in the advertisement, regardless of whether it was the junior or senior synonym. For objective synonyms, we used the same year of description for both names. Continuous variable. This lag value was log-transformed using the formula log(Δ+1) for data normalization.

**Last Publication’s authorship in the WSC**

Type of authorship of the last publication cataloged for the species in the World Spider Catalog (WSC).

The WSC is a reference database for spider taxonomy, providing listings of all currently valid species, genera, and families of spiders, all their synonyms, and all significant taxonomic references to each species as well as their geographic distribution (see example in <https://wsc.nmbe.ch/species/37186>). It is used as the taxonomy backbone for relevant conservation tools such as CITES and GBIF. We evaluated the type of authorship of the last publication cataloged for the species and classified it as being authored by a professional arachnologist or non-professional arachnologist. Non professional authors are defined as authors without formal scientific training and not associated with any academic institution who publish in magazines or journals with no peer-review process and related to the tarantula hobby. Binary variable: non professional or professional.

**Last Publication recorded in the World Spider Catalog**

The lag (Δ) represents the time difference between the year of data collection (2018) and the year of the last paper published about a species, as recorded in the WSC v19.0.

This metric may serve as an indicator of the scientific attention given to a species. A shorter lag indicates recent publications, which suggests that the species has likely undergone taxonomic updates and its identity is well-established. Continuous variable. To ensure data normalization, the lag value was log-transformed using the formula log(Δ+1).

2. **Advertisement Traits**

**Presence of “specifiers” along with the taxa’s name in the advertisement**

Presence of “specifier” words along with the taxa’s name in the advertisement, giving some specificity to the taxon advertised.

Specifiers can be associated with names at species and genus level (e.g., ‘typical color form (TCF)’ in ‘*Pterinochilus murinus* typical color form (TCF)’ and ‘Boa Vista’ in ‘*Avicularia* sp. Boa Vista’). At the species level, it can indicate a variety and at genus level it can indicate a potential species, undescribed or not. Specifiers usually express attractive traits such as color, size, or locality. Binary variable: present or absent.

**Presence of popular name in the advertisement**

Presence of popular names in the advertisement.

Common names are often more accessible and convenient for users to comprehend and search for in online forums, webpages, or web stores compared to scientific names. This accessibility factor can greatly assist potential buyers in locating the specific species of interest. Moreover, common names have the potential to augment the appeal of a species as they tend to highlight distinctive traits, such as the "Eastern horned baboon,"Trinidad dwarf tiger," or "Salmon pink bird-eater." Binary variable: present or absent.

**Size of the specimen advertised**

Leg span of the specimen advertised, in cm.

In cases where multiple sizes are advertised or a size range is provided, the mean value of all the advertised sizes was considered for analysis. Continuous variable.

**Sex of the specimen advertised**

Sex of the specimens advertised.

The specimens advertised were classified into four categories: male, female, couple, or not specified. In terms of this variable, we grouped males and specimens that were not specified into one category, while females and couples (which ultimately include a female) were grouped into another category. This categorization was based on the understanding that females tend to reach maturity later, have larger body sizes, and have longer lifespans than males (Montes de Oca & Mendoza, 2020). Binary variable: female or non-female.

**Life stage of the specimen advertised**

Life stage of the specimen advertised.

We took into account the stated life stage as indicated in the advertisement. If the life stage was not specified, we classified individuals measuring up to 2 inches as spiderlings, individuals nearing the average size of adults for the species as adults, and individuals falling between these two categories as juveniles. Only when the advertisement explicitly mentioned the subadult stage did we consider individuals as such. Information regarding the average size of adults primarily sourced from accessory texts published in the ads, or information in the species description. If the information was lacking in these sources, we searched in other web markets, and in breeders’ or enthusiasts’ blogs, or other online sources (e.g., Arachnoboards, The Tarantula Collective). Although the information sourced from enthusiasts may lack scientific validation and is primarily based on experiential knowledge, it remains relevant for our study objectives as it represents the average size of adult specimens as perceived by the tarantula enthusiasts. Categorical variable: spiderling / juvenile / subadult / adult / not specified.

**Photo of the species in the advertisement**

Presence of photo of the species in the advertisement.

Binary variable: present or absent.

**Source of the specimen advertised**

Source of the specimen advertised.

The source of specimens advertised were classified in captive bred, wild caught, and not specified. As the harvesting from nature to meet the trade can directly impact the wild populations, we grouped the other categories (captive bred and not specified) in one. Binary variable: wild caught or non-wild caught.

**Website**

Website where the advertisement was collected.

Qualitative variable.

**Country of exporting**

Country from where the specimen(s) advertised is shipped from. It is related to the website.

Qualitative character.

**APPENDIX S2** - *Mental Model*

Mental models are cognitive structures that reflect implicit or explicit assumptions of an individual in a particular situation (van der Broek et al. 2021). These cognitive maps indicate relationships among variables, the strength of impact of these elements, helping to understand the dynamics of a system (Gray et al. 2013). Thus, we created a mental model framework (Fig. X) based on Fuzzy-logic Cognitive Mapping using the tool Mental Modeler (<https://www.mentalmodeler.com/>) to express the hypothesized relationship among the variables to be used as our hypothesis in Structural Equation Modeling. See Appendix for further information.

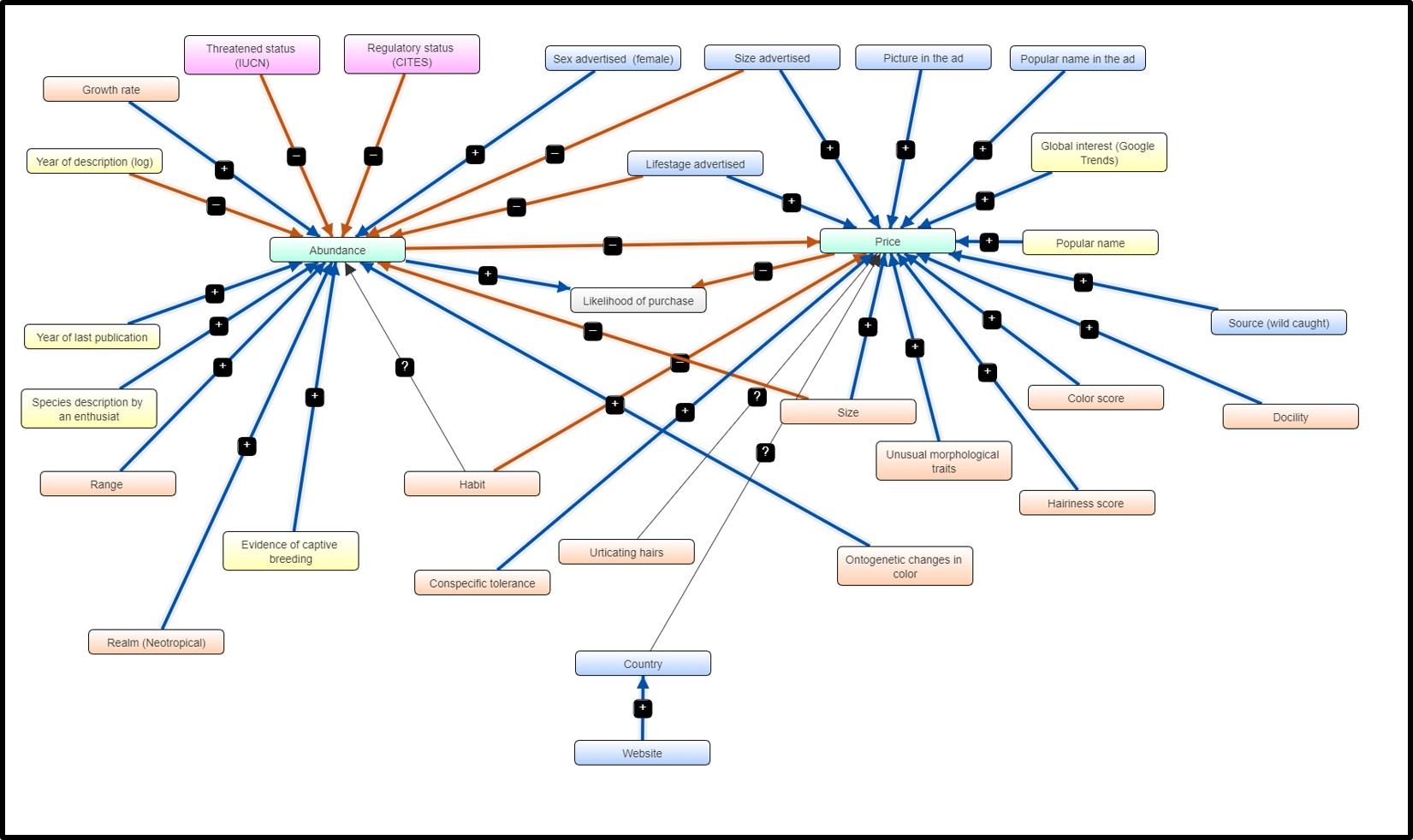

Fig. S2. Mental Model depicting expert hypotheses on relevant traits in the tarantula trade. Boxes: orange = intrinsic traits, yellow = extrinsic aspects, pink = traits related to policy, blue = traits related to the individual advertised.

**APPENDIX 3 -** *Q-Q Plots*

Diagnostic plots for satisfaction of linear regression assumptions for (**A**) species-level and (**B**) advertisement-level models, with log-transformed price as the response variable. Clockwise from top-left: Q-Q plot displaying normality of residuals; residual vs. fitted plot displaying linearity of residuals; Scale-Location plot displaying homoscedasticity; Residuals vs. Leverage plot displaying all points within Cook’s distance of acceptable influence.

Figure S3.A:

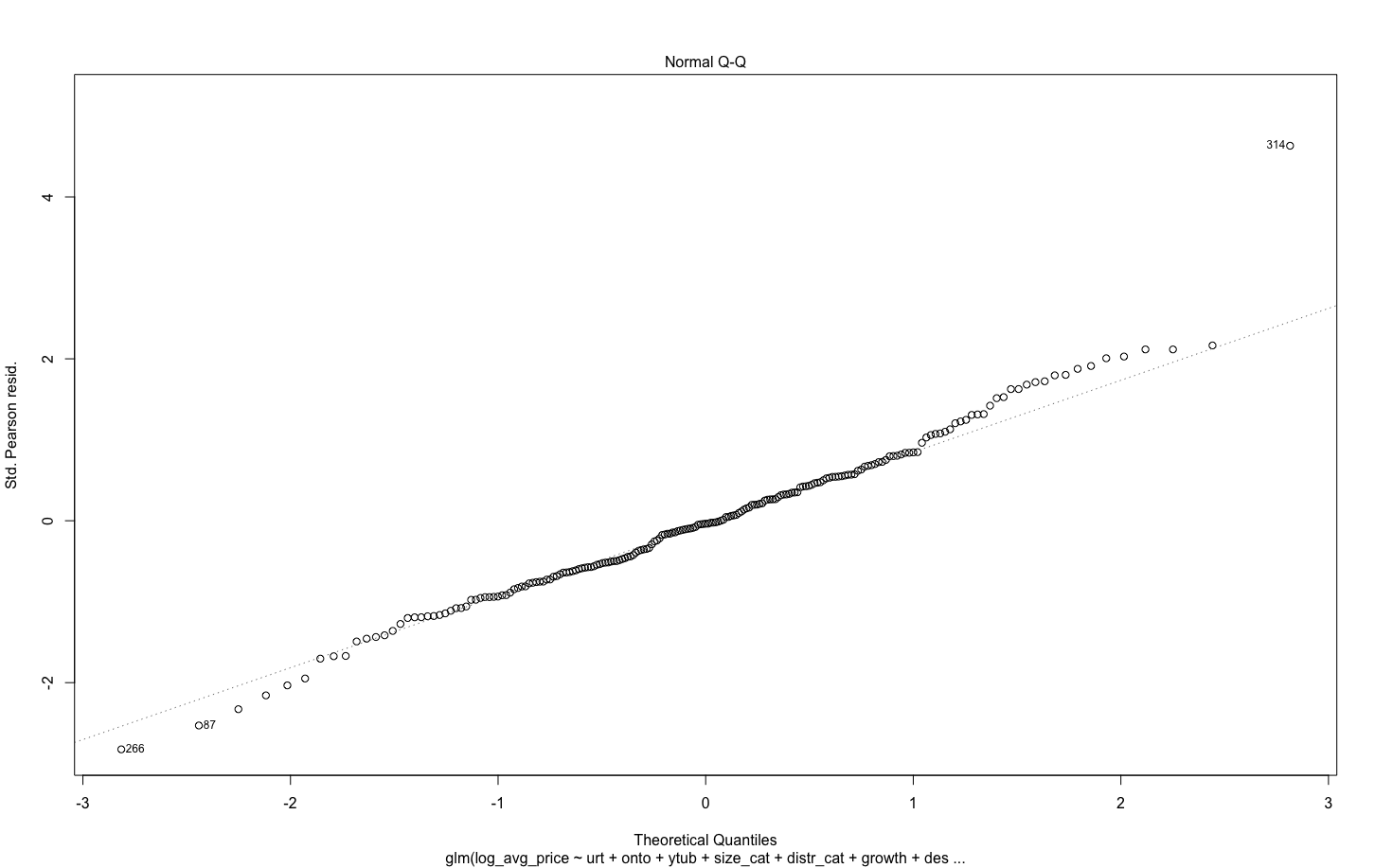

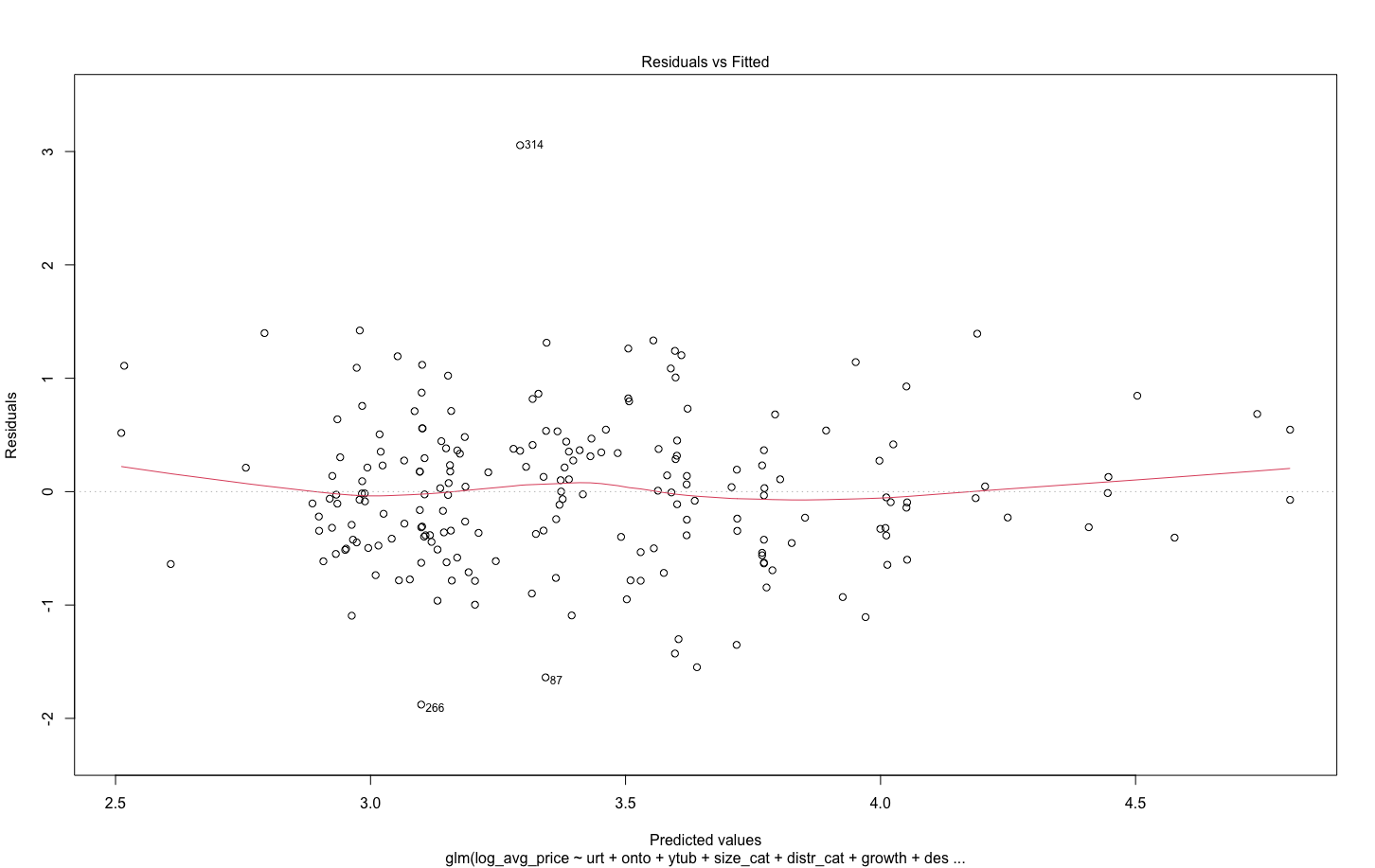

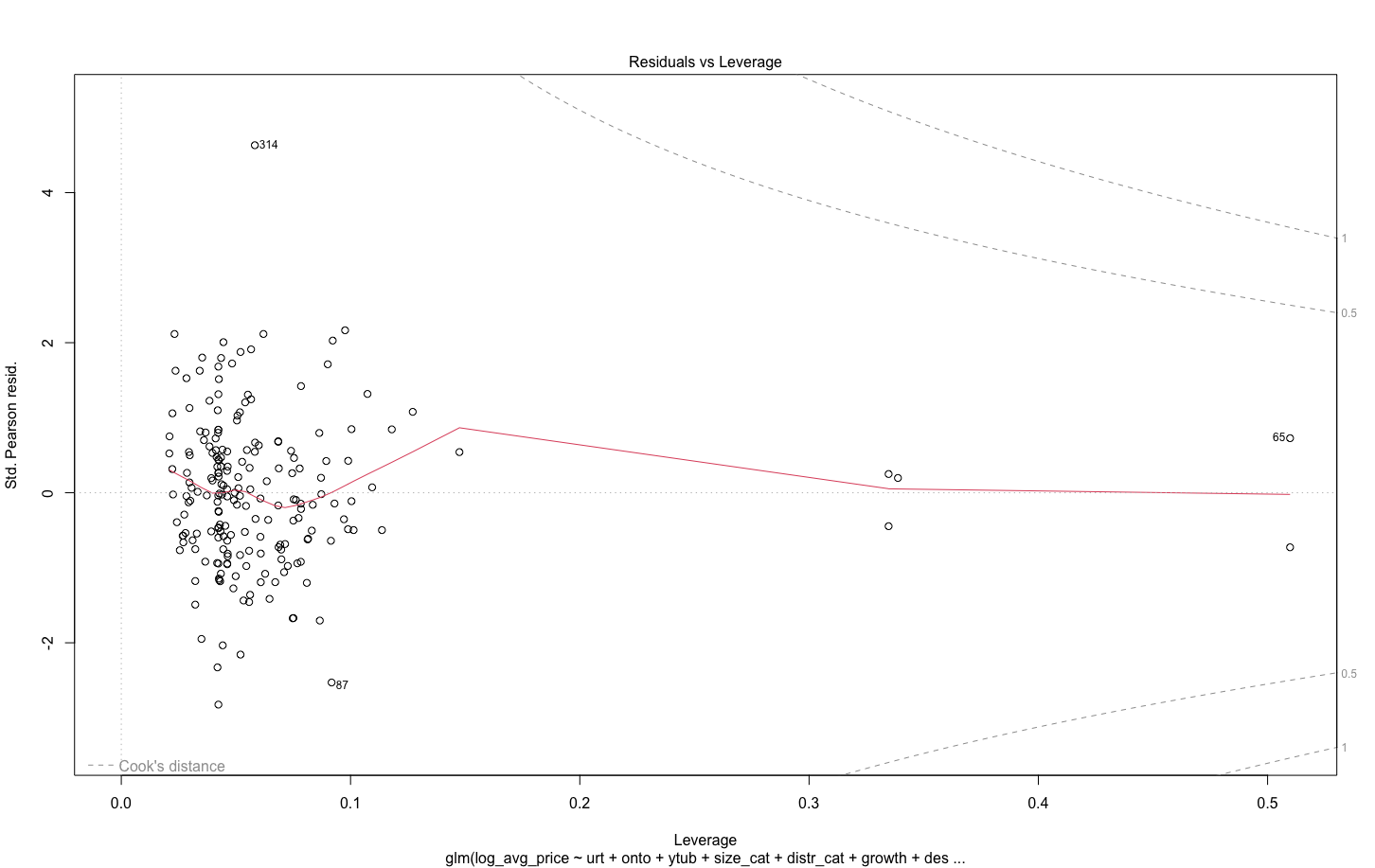

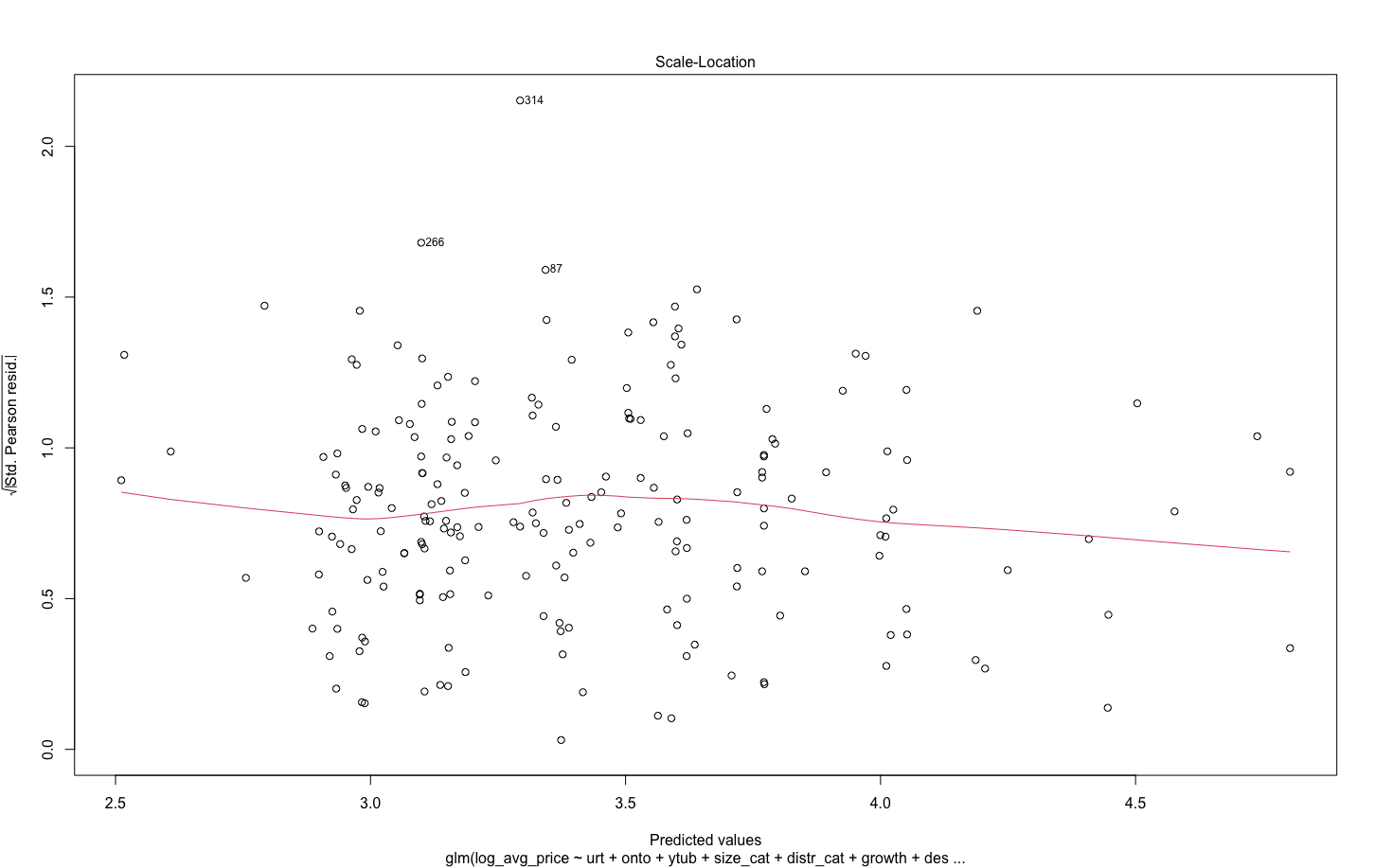

Figure S3.B:

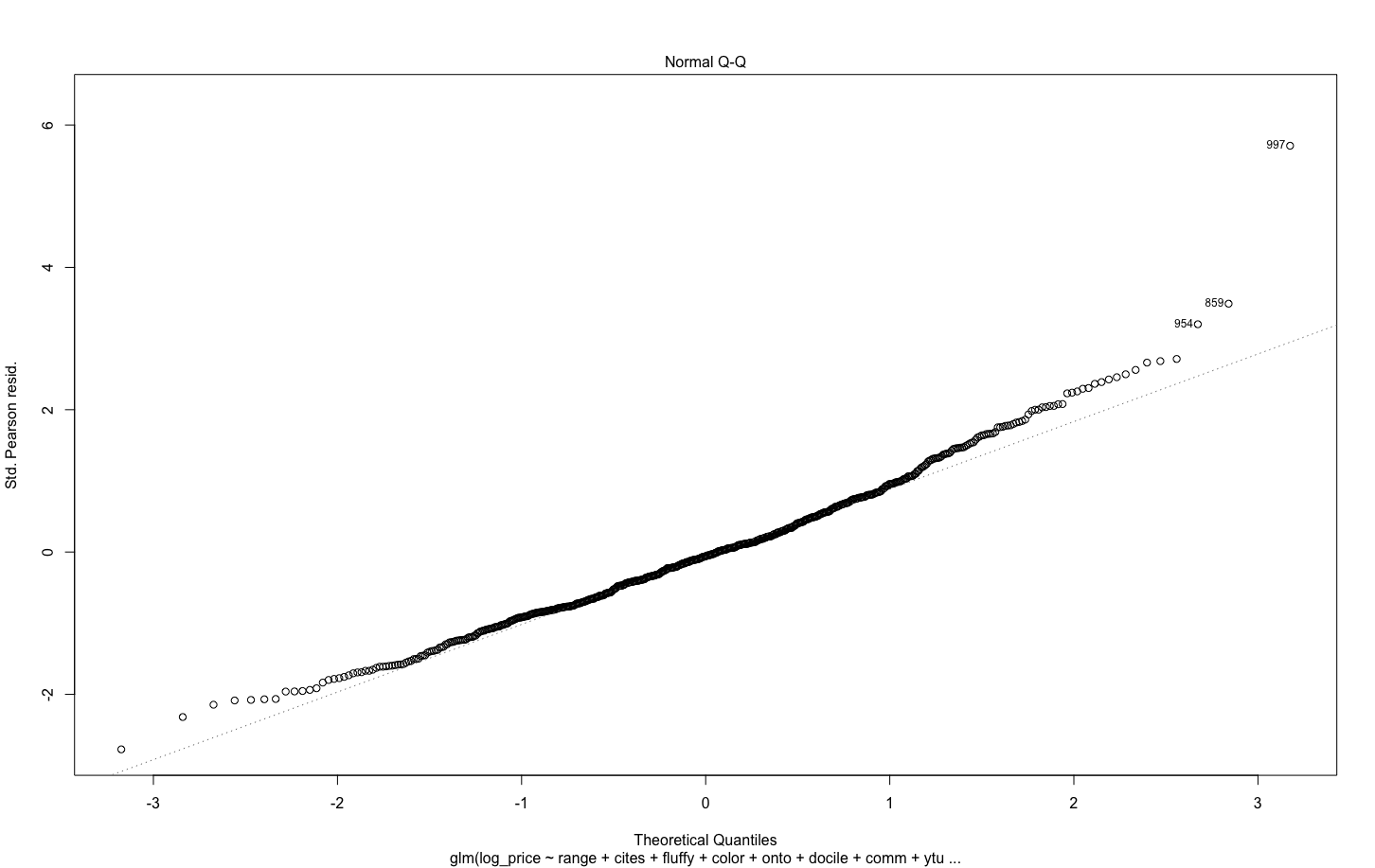

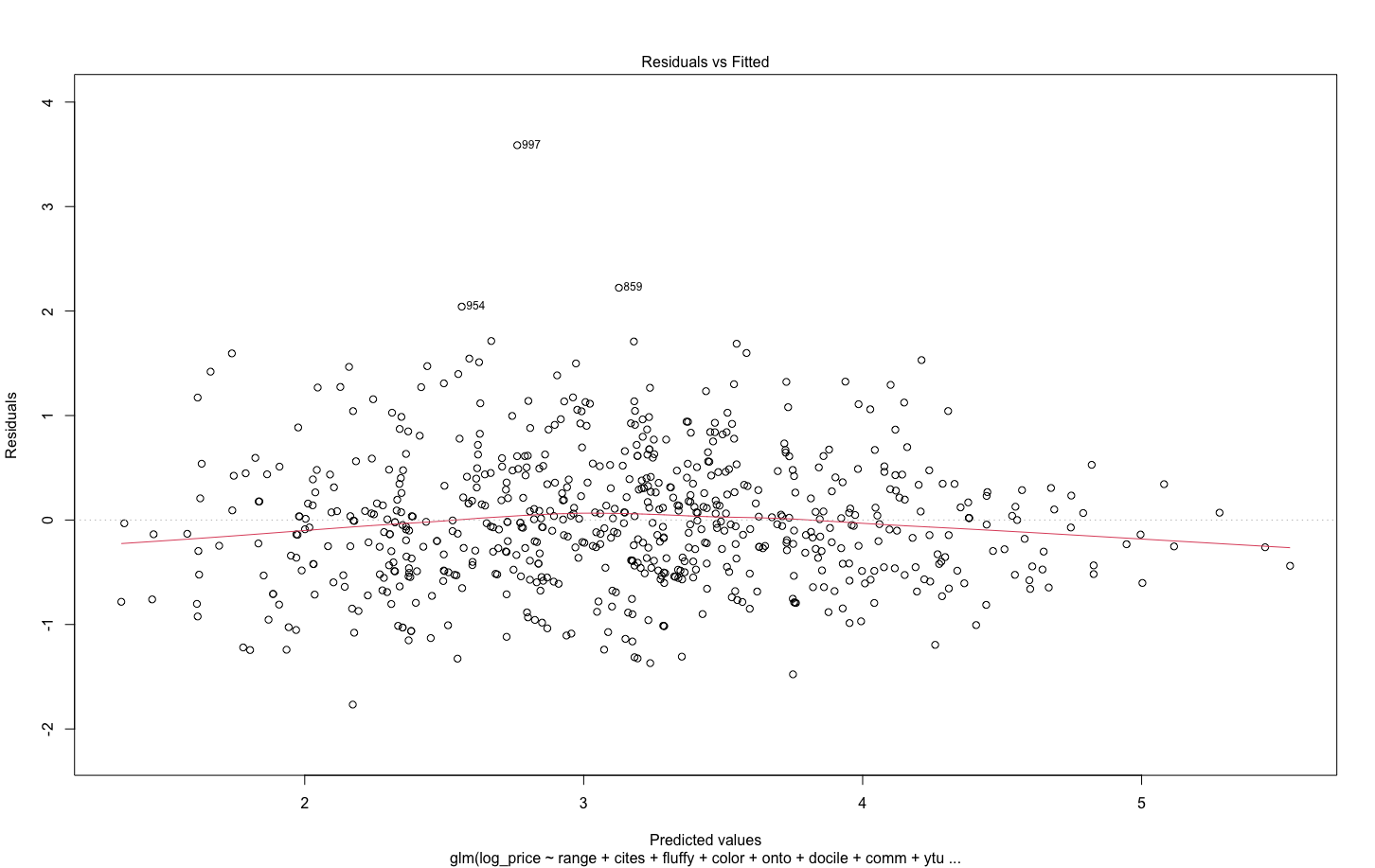

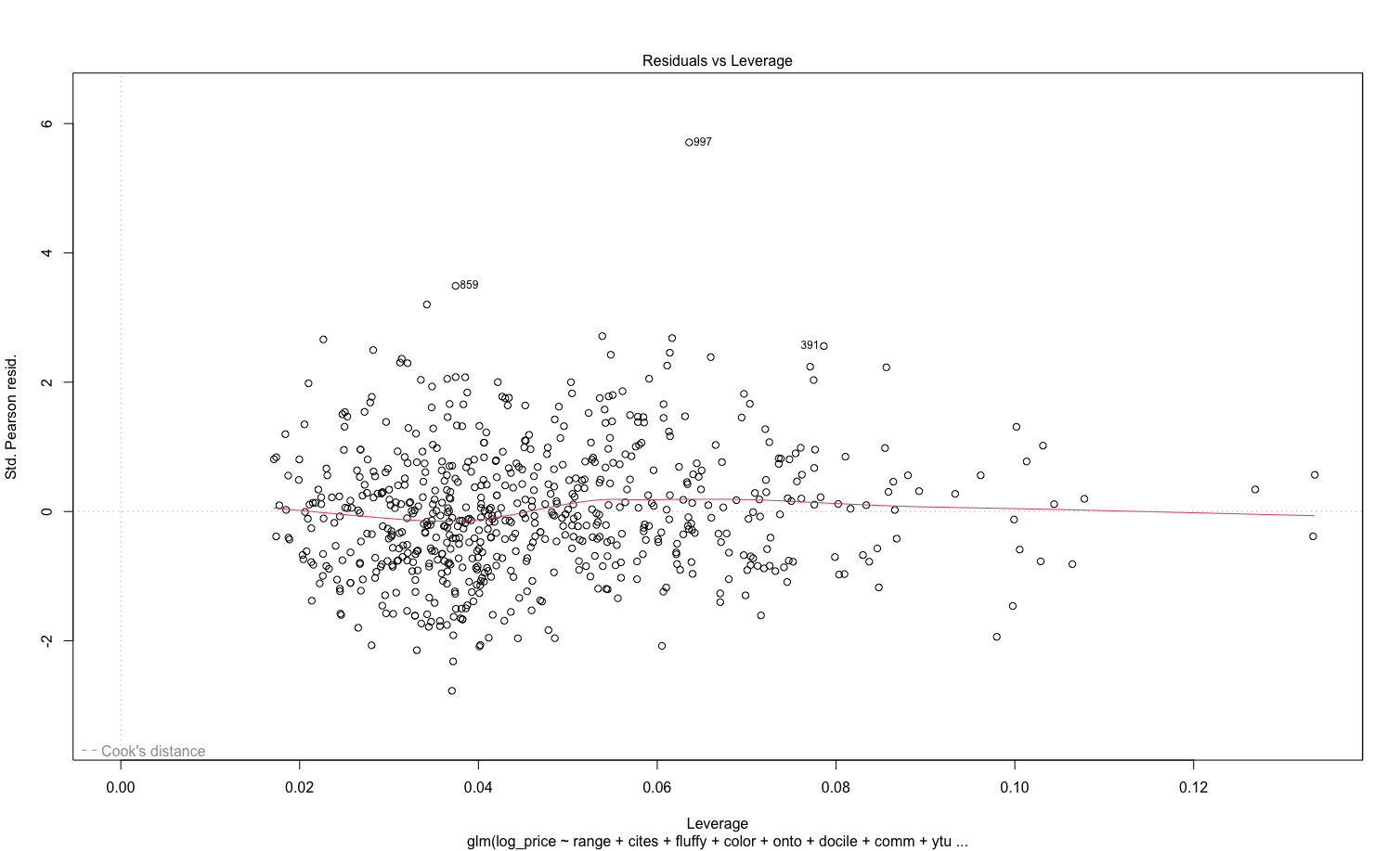

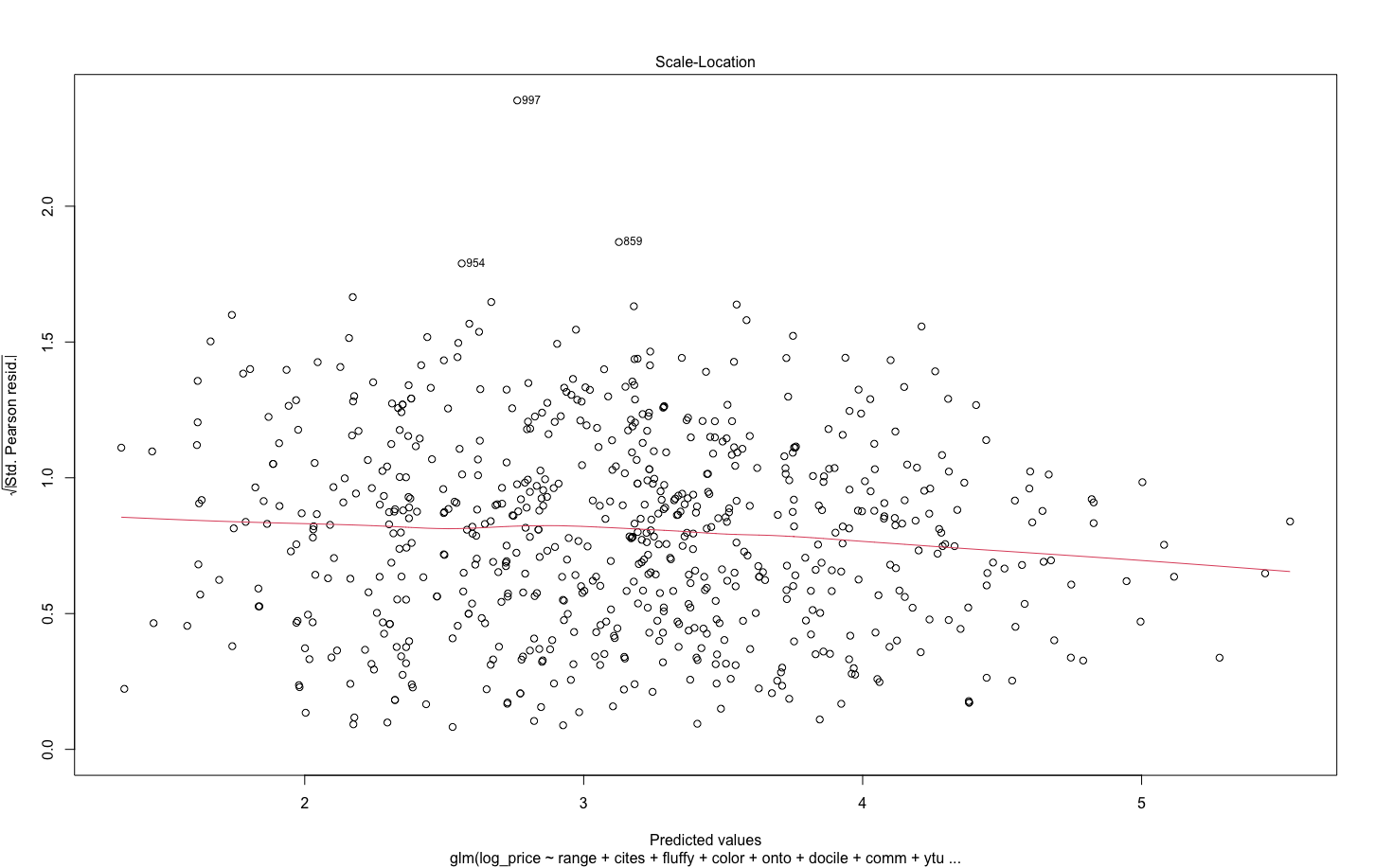

**APPENDIX 4** - *SEM analyses: dSep Tests and Initial Paths*

*dSep Tests*

We performed dSep tests on the initial path and added all variables for which there were significant test results. We repeated this process until there was no further evidence for missing variable paths and we obtained a non-significant Fischer’s C test. If any included variable paths were non-significant, we removed paths in descending order of P-value to determine whether this improved overall model performance (as determined by SEM AIC and Fischer’s C values). To account for sensitivity in our methodology, we repeated the dSep tests on our initial path diagram and added variables in order of dSep test significance, repeating the tests every time a new variable was included. We also repeated both processes for a version of our initial path diagram with the non-significant paths removed. If these multiple approaches resulted in multiple final SEMs with non-significant Fischer’s C tests, we selected the final SEM path diagram with the lowest AIC score. For the species-level SEM, price and abundance were both response variables and therefore dSep tests weare automatically conducted to test for direct relationships between them. In the advertisement-level SEMs, we specifically included market abundance of the advertised species as a variable for consideration in dSep tests, even though abundance itself was not treated as a response.

*Initial Paths*

Table S4.1. Initial SEM path, standard errors and significance values for Species trait Abundance Model, and Species trait Average Price (log transformed) Model SEMs. Covariation between variables is indicated with ‘~’. P-values <0.0005 are denoted with ‘***’, P-values <0.001 are denoted with ‘**’ and P-values <0.05 are denoted with ‘*’. *R* ^2^ Abundance =0.56. *R* ^2^ Average price (log transformed) = 0.23.

| **Response** | **Predictor** | **Estimate** | **Standard Error** | **Degrees of Freedom** | **Critical Value** | **P-value** | **Standard**  **Estimate** |
| --- | --- | --- | --- | --- | --- | --- | --- |
| Abundance | Global interest | 0.8686 | 0.0735 | 194 | 11.8185 | 0*** | - |
| Abundance | Growth rate | -0.1806 | 0.0767 | 194 | -2.3538 | 0.0186* | - |
| Abundance | Hairiness score | 0.2382 | 0.0717 | 194 | 3.3207 | 0.0009*** | - |
| Abundance | Conspecific tolerance | - | - | 1 | 0.0078 | 0.9297 | - |
|  | tolerant | 1.5318 | 0.2069 | Inf | 7.4047 | 0*** | - |
|  | not tolerant | 1.5521 | 0.0724 | Inf | 21.4336 | 0*** | - |
| Abundance | Docility | - | - | 1 | 11.5623 | 0.0008*** | - |
|  | docile | 1.2744 | 0.1447 | Inf | 8.8047 | 0*** | - |
|  | non docile | 1.8095 | 0.1217 | Inf | 14.8672 | 0*** | - |
| Abundance | Last publication’s authorship | - | - | 1 | 7.0408 | 0.0086** | - |
|  | non professional | 1.3531 | 0.1387 | Inf | 9.7534 | 0*** | - |
|  | professional | 1.7308 | 0.12 | Inf | 14.4286 | 0*** | - |
| Average Price (log) | Growth rate | 0.0297 | 0.0672 | 195 | 0.4428 | 0.6584 | - |
| Average Price (log) | Year of description (log) | -0.141 | 0.0481 | 195 | -2.9325 | 0.0038** | - |
| Average Price (log) | Size | - | - | 2 | 8.1198 | 0.0004*** | - |
|  | small | 3.4582 | 0.1807 | 195 | 19.1344 | 0*** | - |
|  | medium | 3.5533 | 0.1178 | 195 | 30.1523 | 0*** | - |
|  | large | 4.1073 | 0.1816 | 195 | 22.6169 | 0*** | - |
| Average Price (log) | Distribution category | - | - | 5 | 5.3984 | 0.0001*** | - |
|  | Afrotropical | 3.3656 | 0.1813 | 195 | 18.5611 | 0*** | - |
|  | Neotropical | 3.3912 | 0.0706 | 195 | 48.0578 | 0*** | - |
|  | Indo-Malayan | 3.4379 | 0.1322 | 195 | 26.0004 | 0*** | - |
|  | Palearctic | 3.7181 | 0.5079 | 195 | 7.3199 | 0*** | - |
|  | Australasian | 3.8644 | 0.4252 | 195 | 9.0885 | 0*** | - |
|  | Nearctic | 4.4606 | 0.2069 | 195 | 21.5565 | 0*** | - |
| Urticating Hairs | ~~Distribution | 0.7016 | - | 224 | 14.7371 | 0*** | 0.7016 |
| Docility | ~~Urticating Hairs | 0.5319 | - | 211 | 9.1241 | 0*** | 0.5319 |
| Docility | ~~Distribution | 0.3659 | - | 211 | 5.7109 | 0*** | 0.3659 |

Tab. S4.2 - Initial SEM path, standard errors and significance values for Advertisement trait model, variable response “price” (log transformed) SEM. Covariation between variables is indicated with ‘~’. P-values <0.0005 are denoted with ‘***’, P-values <0.001 are denoted with ‘**’ and P-values <0.05 are denoted with ‘*’. For the predictors marked with ^a^ , the response variable is “urticating hairs”, and for ^b^ the response variable is “docility”. Country codes for the web stores: CAN = Canada, GER = Germany, POL = Poland, SA = South Africa, UK = United Kingdom, USA = United States of America. *R* ^2^ =.0.45

| **Response** | **Predictor** | **Estimate** | **Standard Error** | **Degrees of Freedom** | **Critical Value** | **P-value** | **Standard**  **Estimate** |
| --- | --- | --- | --- | --- | --- | --- | --- |
| Price (log) | Size advertised | -6.00E-04 | 0.001 | 604 | -0.6332 | 0.5268 | - |
| Price (log) | Growth rate | 0.0043 | 0.0298 | 604 | 0.1427 | 0.8866 | - |
| Price (log) | Sex | - | - | 1 | 40.0233 | 0*** | - |
|  | non female | 3.348 | 0.1026 | 604 | 32.6272 | 0*** | - |
|  | female | 3.9557 | 0.1026 | 604 | 38.5662 | 0*** | - |
| Price (log) | Life stage advertised | - | - | 3 | 23.5619 | 0*** | - |
|  | spiderling | 3.1094 | 0.0642 | 604 | 48.4645 | 0*** | - |
|  | juvenile | 3.6425 | 0.0541 | 604 | 67.3053 | 0*** | - |
|  | adult | 3.6915 | 0.3142 | 604 | 11.7499 | 0*** | - |
|  | subadult | 4.164 | 0.1592 | 604 | 26.1512 | 0*** | - |
| Price (log) | Website | - | - | 7 | 49.3719 | 0*** | - |
|  | POL2 | 2.609 | 0.1413 | 604 | 18.4664 | 0*** | - |
|  | POL1 | 3.1688 | 0.1099 | 604 | 28.8298 | 0*** | - |
|  | SA | 3.2569 | 0.1238 | 604 | 26.3094 | 0*** | - |
|  | GER | 3.5142 | 0.1787 | 604 | 19.6646 | 0*** | - |
|  | UK | 3.5197 | 0.1481 | 604 | 23.761 | 0*** | - |
|  | CAN | 3.7587 | 0.1227 | 604 | 30.6451 | 0*** | - |
|  | USA1 | 4.6341 | 0.1425 | 604 | 32.5259 | 0*** | - |
|  | USA2 | 4.7533 | 0.1529 | 604 | 31.0875 | 0*** | - |
| Price (log) | Popular name in the ad | - | - | 1 | 30.1111 | 0*** | - |
|  | present | 3.3282 | 0.1016 | 604 | 32.7557 | 0*** | - |
|  | absent | 3.9754 | 0.1143 | 604 | 34.7769 | 0*** | - |
| Urticating hairs | ~~Distribution | 0.6773 | - | 707 | 24.4767 | 0*** | 0.6773 |
| Docility | ~~Urticating hairs | 0.4944 | - | 686 | 14.8989 | 0*** | 0.4944 |
| Docility | ~~Distribution | 0.2808 | - | 686 | 7.6618 | 0*** | 0.2808 |

Tab. S4.3 - Initial SEM path, standard errors and significance values for Advertisement trait model, Mixed Model, variable response “price” (log transformed) SEM. Covariation between variables is indicated with ‘~’. P-values <0.0005 are denoted with ‘***’, P-values <0.001 are denoted with ‘**’ and P-values <0.05 are denoted with ‘*’. For the predictors marked with ^a^ , the response variable is “urticating hairs”, and for ^b^ the response variable is “docility”. *R* ^2^ =.0.55

| **Response** | **Predictor** | **Estimate** | **Standard Error** | **Degrees of Freedom** | **Critical Value** | **P-value** | **Standard**  **Estimate** |
| --- | --- | --- | --- | --- | --- | --- | --- |
| Price (log) | Year of description | -0.121 | 0.0302 | 583.6437 | -4.0119 | 0.0001*** | - |
| Price (log) | Size advertised | -1.00E-03 | 9.00E-04 | 583.2993 | -1.116 | 0.2649 | - |
| Price (log) | Growth rate | -0.0033 | 0.0412 | 583.3214 | -0.0801 | 0.9362 | - |
| Price (log) | Captive breeding evidence | - | - | 1 | 23.1825 | 0*** | - |
|  | present | 4.039 | 0.2621 | 10.5522 | 15.4132 | 0*** | - |
|  | absent | 4.5581 | 0.2784 | 13.4197 | 16.3728 | 0*** | - |
| Price (log) | Tolerance to conspecifics | - | - | 1 | 20.5371 | 0*** | - |
|  | tolerant | 4.0445 | 0.2725 | 12.3285 | 14.8412 | 0*** | - |
|  | not tolerant | 4.5526 | 0.269 | 11.7188 | 16.9218 | 0*** | - |
| Price (log) | Docility | - | - | 1 | 22.5009 | 0*** | - |
|  | non docile | 4.1309 | 0.2631 | 10.7234 | 15.7011 | 0*** | - |
|  | docile | 4.4662 | 0.2714 | 12.124 | 16.4581 | 0*** | - |
| Price (log) | Size category | - | - | 2 | 22.2235 | 0*** | - |
|  | small | 4.0797 | 0.2769 | 13.128 | 14.7339 | 0*** | - |
|  | medium | 4.0942 | 0.2612 | 10.423 | 15.6736 | 0*** | - |
|  | large | 4.7218 | 0.2792 | 13.5709 | 16.9136 | 0*** | - |
| Price (log) | Distribution category | - | - | 5 | 9.3752 | 0*** | - |
|  | Neotropical | 3.6991 | 0.2411 | 7.5783 | 15.3403 | 0*** | - |
|  | Afrotropical | 4.1464 | 0.2638 | 10.8575 | 15.7157 | 0*** | - |
|  | Indo-Malayan | 4.2026 | 0.2573 | 9.8082 | 16.3365 | 0*** | - |
|  | Nearctic | 4.371 | 0.2763 | 13.0222 | 15.8185 | 0*** | - |
|  | Australasian | 4.6067 | 0.3873 | 48.4047 | 11.895 | 0*** | - |
|  | Palearctic | 4.7655 | 0.5592 | 172.2422 | 8.5216 | 0*** | - |
| Price (log) | Sex | - | - | 1 | 42.6646 | 0*** | - |
|  | non-female | 4.0034 | 0.269 | 11.6769 | 14.883 | 0*** | - |
|  | female or couple | 4.5938 | 0.2685 | 11.635 | 17.1089 | 0*** | - |
| Price (log) | Life stage | - | - | 3 | 19.6612 | 0*** | - |
|  | spiderling | 3.7716 | 0.2567 | 9.7227 | 14.6926 | 0*** | - |
|  | juvenile | 4.2402 | 0.2541 | 9.3391 | 16.689 | 0*** | - |
|  | subadult | 4.4545 | 0.3894 | 49.3025 | 11.4398 | 0*** | - |
|  | adult | 4.7279 | 0.29 | 15.7042 | 16.3057 | 0*** | - |
| Price (log) | Ontogenetic changes in color | - | - | 1 | 18.7645 | 0*** | - |
|  | absent | 4.1008 | 0.26 | 10.2274 | 15.77 | 0*** | - |
|  | present | 4.4963 | 0.2773 | 13.2213 | 16.2128 | 0*** | - |
| Urticating hairs | ~~Distribution | 0.6773 | - | 707 | 24.4767 | 0*** | 0.6773 |
| Docility | ~~Urticating Hairs | 0.4944 | - | 686 | 14.8989 | 0*** | 0.4944 |
| Docility | ~~Distribution | 0.2808 | - | 686 | 7.6618 | 0*** | 0.2808 |

**APPENDIX S5 -** *List of Taxa (valid or invalid) found in the ads* *(as in WSC 19.0)*

| **List of taxa present in the ads** | | | | | |
| --- | --- | --- | --- | --- | --- |
| *Acanthoscurria chacoana*  Brèthes 1909 | *Brachypelma hamorii*  Tesmoingt, Cleton & Verdez 1997 | *Cyriopagopus schmidti*  (von Wirth 1991) | *Holothele sanguiniceps*  (F. O. Pickard-Cambridge, 1899) | *Pamphobeteus* sp. | *Pterinochilus lugardi*  Pocock 1900 |
| *Acanthoscurria geniculata*  (C. L. Koch 1841) | *Brachypelma kahlenbergi*  (Rudloff 2008) | *Cyriopagopus* sp. | *Holothele sericea*  (Simon 1903) | *Pelinobius muticus*  Karsch 1885 | *Pterinochilus murinus*  (Walckenaer, 1837) |
| *Acanthoscurria musculosa*  Simon 1892 | *Brachypelma klaasi*  (Schmidt & Krause 1994) | *Cyriopagopus vonwirthi*  Schmidt 1995 | *Holothele* sp. | *Phlogiellus* sp*.* | *Pterinochilus* sp*.* |
| *Acanthoscurria natalensis*  Chamberlin 1917 | *Brachypelma sabulosum*  (F. O. Pickard-Cambridge 1897) | *Cyrtopholis cursor*  (Ausserer 1875) | *Homoeomma* sp*.* | *Phlogius crassipes*  (L. Koch 1874) | *Pterinopelma sazimai*  Bertani, Nagahama & Fukushima 2011 |
| *Acanthoscurria paulensis*  Mello-Leitão 1923 | *Brachypelma schroederi*  (Rudloff 2003) | *Davus pentaloris*  (Simon 1888) | *Hysterocrates gigas*  Pocock 1897 | *Phormictopus atrichomatus*  Schmidt 1991 | *Sahydroaraneus raja*  (Gravely 1915) |
| *Acanthoscurria* sp*.* | *Brachypelma smithi*  (F. O. Pickard-Cambridge 1897) | *Davus ruficeps*  (Simon 1891) | *Hysterocrates Hercules*  Pocock 1900 | *Phormictopus auratus*  Ortiz & Bertani 2005 | *Schizopelma bicarinatum*  F. O. Pickard-Cambridge 1897 |
| *Acanthoscurria theraphosoides*  (Doleschall 1871) | *Brachypelma* sp. | *Dolichothele diamantinensis*  (Bertani, Santos & Righi 2009) | *Hysterocrates laticeps*  Pocock 1897 | *Phormictopus cancerides*  Latreille 1806 | *Selenocosmia arndsti*  (Schmidt & von Wirth 1991) |
| *Ami* sp. | *Brachypelma vagans*  (Ausserer 1875) | *Dolichothele exilis*  Mello-Leitão 1923 | *Hysterocrates sp.* | *Phormictopus cubensis*  Chamberlin 1917 | *Selenocosmia aruana*  Strand 1911 |
| *Aphonopelma anax*  (Chamberlin 1940) | *Brachypelma verdezi*  (Schmidt 2003) | *Encyocratella olivacea*  Stand 1907 | *Idiothele mira*  Gallon 2010 | *Phormictopus* sp. | *Selenocosmia crassipes*  (L. Koch 1874) |
| *Aphonopelma bicoloratum*  Struchen, Brändle & Schmidt 1996 | *Bumba cabocla*  (Pérez-Miles 2000) | *Ephebopus cyanognathus*  West & Marshall 2000 | *Kochiana brunnipes*  (C. L. Koch 1842) | *Phormingochilus carpenter*  (Smith & Jacobi 2015) | *Selenocosmia kovariki*  (Schmidt & Krause 1995) |
| *Aphonopelma chalcodes*  Chamberlin 1940 | *Caribena laeta*  (C. L. Koch 1842) | *Ephebopus murinus*  (Walckenaer 1837) | *Lampropelma nigerrimum*  Simon 1892 | *Phormingochilus everetti*  Pocok 1895 | *Selenocosmia* sp. |
| *Aphonopelma crinirufum*  (Valerio 1980) | *Caribena versicolor*  (Walckenaer 1837) | *Ephebopus rufescens*  West & Marshall 2000 | *Lampropelma nigerrimum arboricola*  Schmidt & Barensteiner 2015 | *Phormingochilus* sp*.* | *Selenocosmiinae* sp. |
| *Aphonopelma eutylenum*  Chamberlin 1940 | *Catumiri argentinensis*  (Mello-Leitão 1941) | *Ephebopus uatuman*  Lucas, Silva & Bertani 1992 | *Lampropelma* sp. | *Phrixotrichus* sp. | *Sericopelma generala*  Valerio 1980 |
| *Aphonopelma gabeli*  Smith 1995 | *Catumiri parvum*  (Keyserling 1878) | *Euathlus parvulus*  (Pocock 1903) | *Lampropelma violaceopes*  Abraham 1924 | *Phrixotrichus vulpinus*  Simon 1889 | *Sericopelma melanotarsum*  Valerio 1980 |
| *Aphonopelma hentzi*  (Girard 1852) | *Ceratogyrus darling*  Pocock 1897 | *Euathlus pulcherrimaklaasi*  (Schmidt 1991) | *Lasiodora difficilis*  Mello-Leitão 1921 | *Plesiopelma longisternale*  (Schiapelli & Gerschman 1942) | *Sericopelma rubronitens*  Ausserer 1875 |
| *Aphonopelma iodius*  (Chamberlin & Ivie 1939) | *Ceratogyrus marshalli*  Pocock 1897 | *Euathlus sp.* | *Lasiodora klugi*  (C. L. Koch 1841) | *Poecilotheria fasciata*  Latreille 1804 | *Sericopelma* sp*.* |
| *Aphonopelma johnnycashi*  Hamilton 2016 | *Ceratogyrus meridionalis*  (Hirst 1907) | *Eupalaestrus campestratus*  (Simon 1891) | *Lasiodora parahybana*  Mello-Leitão 1971 | *Poecilotheria formosa*  Pocock 1899 | *Sphaerobothria hoffmanni*  Karsch 1879 |
| *Aphonopelma mareki*  Hamilton, Hendrixson & Bond 2016 | *Ceratogyrus sanderi*  Strand 1906 | *Euthycaelus colonica*  (Simon 1889) | *Lasiodorides polycuspulatus*  Schmidt & Bischoff 1997 | *Poecilotheria hanumavilasumica*  Smith 2004 | *Stichoplastoris asterix*  (Valerio 1980) |
| *Aphonopelma marxi*  (Simon 1891) | *Chaetopelma olivaceum*  (C. L. Koch 1841) | *Grammostola actaeon*  Pocock 1903 | *Megaphobema mesomelas*  (O. Pickard-Cambridge 1892) | *Poecilotheria metallica*  Pocock 1899 | *Stromatopelma calceatum*  (Fabricius 1793) |
| *Aphonopelma moderatum*  (Chamberlin & Ivie) 1939 | *Chilobrachys andersoni*  (Pocock 1895) | *Grammostola grossa*  Ausserer 1871 | *Megaphobema robustum*  Ausserer 1875 | *Poecilotheria Miranda*  Pocock 1900 | *Tapinauchenius cupreus*  Schmidt & Bauer 1996 |
| *Aphonopelma Paloma*  Prentice 1993 | *Chilobrachys dyscolus*  (Simon 1886) | *Grammostola iheringi*  Keyserling 1891 | *Megaphobema velvetosoma*  Schmidt 1995 | *Poecilotheria ornate*  Pocock 1899 | *Tapinauchenius gigas*  (Caporiacco 1954) |
| *Aphonopelma seemanni*  (F. O. Pickard-Cambridge 1897) | *Chilobrachys fimbriatus*  Pocock 1899 | *Grammostola porter*  Mello-Leitão 1936 | *Metriopelma familiar*  (Simon 1889) | *Poecilotheria regalis*  Pocock 1899 | *Tapinauchenius* sp*.* |
| *Aphonopelma superstitionense*  Hamilton, Hendrixson & Bond 2016 | *Chilobrachys fumosus*  (Pocock 1895) | *Grammostola pulchra*  Mello-Leitão 1921 | *Monocentropus balfouri*  Pocock 1897 | *Poecilotheria rufilata*  Pocock 1899 | *Tapinauchenius violaceus*  (Mello-Leitão 1930) |
| *Aphonopelma vorhiesi*  (Chamberlin & Ivie 1939) | *Chilobrachys huahini*  Schmidt & Huber 1996 | *Grammostola pulchripes*  Simon 1891 | *Neoholothele incei*  (F. O. Pickard-Cambridge 1899) | *Poecilotheria smithi*  Kirk 1996 | *Theraphosa apophysis*  (Tinter 1991) |
| *Aphonopelma xwalxwal*  Hamilton 2016 | *Chilobrachys paviei*  (Simon 1886) | *Grammostola rosea*  Walckenaer 1837 | *Neostenotarsus* sp. | *Poecilotheria* sp. | *Theraphosa blondi*  Latreille 1804 |
| *Augacephalus ezendami*  (Gallon 2001) | *Chilobrachys* sp. | *Hapalopus formosus*  Ausserer 1875 | *Nhandu carapoensis*  Lucas 1983 | *Poecilotheria striata*  Pocock 1895 | *Theraphosa stirmi*  Rudloff & Weinmann 2010 |
| *Avicularia aurantiaca*  Bauer 1996 | *Chromatopelma cyaneopubescens*  (Strand 1907) | *Hapalopus* sp*.* | *Nhandu chromatus*  Schmidt 2004 | *Poecilotheria subfusca*  Pocock 1895 | *Theraphosidae* sp*.* |
| *Avicularia avicularia*  (Linnaeus 1758) | *Citharacanthus cyaneus*  (Rudloff 994) | *Hapalopus triseriatus*  Caporiacco 1955 | *Nhandu coloratovillosus*  Schmidt 1998 | *Poecilotheria tigrinawesseli*  Smith 2006 | *Theraphosinae* sp. |
| *Avicularia braunshauseni*  Tesmoingt 1999 | *Citharognathus tongmianensis*  Zhu, Li & Song 2002 | *Haploclastus devamatha*  Prasanth & Sunil Jose 2014 | *Nhandu* sp. | *Poecilotheria vittata*  Pocock 1895 | *Thrigmopoeus psychedelicus*  Sanap & Mirza 2014 |
| *Avicularia juruensis*  Mello-Leitão 1923 | *Coremiocnemis cunicularia*  (Simon 1892) | *Haplocosmia himalayana*  (Pocock 1899) | *Nhandu tripepii*  (Dresco 1984) | *Proshapalopus amazonicus*  Bertani 2001 | *Thrigmopoeus truculentus*  Pocock 1899 |
| *Avicularia merianae*  Fukushima & Bertani 2017 | *Coremiocnemis hoggi*  West & Nunn 2010 | *Haplocosmia nepalensis*  Schmidt & von Wirth 1996 | *Omothymus schioedtei*  Thorell 1891 | *Psalmopoeus cambridgei*  Pocock 1895 | *Thrixopelma ockerti*  Schmidt 1994 |
| *Avicularia metallica*  (Ausserer 1875) | *Coremiocnemis obscura*  West & Nunn 2010 | *Haplocosmia sp.* | *Ornithoctoninae* sp. | *Psalmopoeus ecclesiasticus*  Pocock 1903 | *Thrixopelma pruriens*  Schmidt 1998 |
| *Avicularia minatrix*  Pocock 1903 | *Cyclosternum fasciatus*  (Mello-Leitão 1930) | *Haplopelma sp.* | *Ornithoctonus aureotibialis*  von Wirth & Striffler 2005 | *Psalmopoeus irminia*  Saager 1994 | *Thrixopelma* sp. |
| *Avicularia purpurea*  Kirk 1990 | *Cyclosternum* sp. | *Harpactira baviana*  Purcell 1902 | *Ornithoctonus* sp. | *Psalmopoeus langenbucheri*  Schmidt, Bullmer & Thierer-Lutz 2006 | *Vitalius paranaensis*  Bertani 2001 |
| *Avicularia rufa*  Schiapelli & Gerschman 1945 | *Cyriocosmus aueri*  Kaderka 2016 | *Harpactira cafreriana*  Walckenaer 1837 | *Orphnaecus dichromatus*  (Schmidt & von Wirth 1992) | *Psalmopoeus pulcher*  Petrunkevitch 1925 | *Xenesthis immanis*  Ausserer 1875 |
| *Avicularia sooretama*  (Bertani & Fukushima 2009) | *Cyriocosmus chicoi*  Pérez-Milles 1998 | *Harpactira ditactor*  Purcell 1902 | *Orphnaecus philippinus*  Schmidt 1999 | *Psalmopoeus reduncus*  (Karsch 1880) | *Xenesthis* sp. |
| *Avicularia* sp. | *Cyriocosmus elegans*  (Simon 1889) | *Harpactira marksi*  Purcell 1902 | *Orphnaecus* sp*.* | *Psalmopoeus victori*  Mendoza 2014 | *Ybyrapora diversipes*  (C. L. Koch 1842) |
| *Bonnetina cyaneifemur*  Vol 2000 | *Cyriocosmus leetzi*  Vol 1999 | *Harpactira pulchripes*  Pocock 1901 | *Pachistopelma bromelicola*  Bertani 2012 | *Psednocnemis brachyramosa*  (West & Nunn 2010) | *Ybyrapora sooretama*  (Bertani & Fukushima 2009) |
| *Brachypelma albiceps*  Pocock 1903 | *Cyriocosmus perezmilesi*  Kardeka 2007 | *Harpactirella lightfooti*  Purcell 1902 | *Pamphobeteus antinous*  Pocock 1903 | *Psednocnemis davidgohi*  West, Nunn & Hogg 2012 | [*Haplopelma robustum*](http://wsc.nmbe.ch/species/37473/Cyriopagopus_robustus)  Strand 1907 |
| *Brachypelma albopilosum*  Valerio 1980 | *Cyriocosmus ritae*  Pérez-Milles 1998 | *Harpactirella overdijki*  Gallon 2010 | *Pamphobeteus fortis*  Ausserer 1875 | *Psednocnemis jeremyhuffi*  (West & Nunn 2010) | [*Haplopelma schmidti*](http://wsc.nmbe.ch/species/37473/Cyriopagopus_robustus)  von Wirth 1991 |
| *Brachypelma auratum*  Schmidt 1992 | *Cyriopagopus hainanus*  (Liang, Peng, Huang & Chen 999) | *Heteroscodra maculate*  Pocock 1900 | *Pamphobeteus grandis*  Bertani, Fukushima & Silva 2008 | *Pseudhapalopus* sp*.* | *Typhochlaena seladonia*  (C. L. Koch 1841) |
| *Brachypelma baumgarteni*  Smith 1993 | *Cyriopagopus lividus*  Smith 1996 | *Heterothele gabonensis*  Lucas 1858 | *Pamphobeteus insignis*  Pocock 1903 | *Pseudhapalopus trinitatis*  (Pocock 1903) |  |
| *Brachypelma boehmei*  Schmidt & Klaas 1993 | *Cyriopagopus longipes*  (von Wirth & Striffler 2005) | *Heterothele villosella*  Strand 1907 | *Pamphobeteus nigricolor*  Ausserer 1875 | *Pterinochilus chordates*  (Gerstäcker 1873) |  |
| *Brachypelma emilia*  (White 1856) | *Cyriopagopus minax*  Thorell 1897 | *Holothele longipes*  (L. Koch 1875) | *Pamphobeteus petersi*  Schmidt 2002 | *Pterinochilus lapalala*  Gallon & Engelbrecht 2011 |  |

**APPENDIX S6 -** *Taxa and ‘trade names’ advertised per websites.*

Table S6.1. Count of ads selling tarantulas based on scientific names and 'trade names' in each webstore, and number of distinct species and 'trade names' found in the ads of each webstore. Country codes for the web stores: CAN = Canada, GER = Germany, POL = Poland, SA = South Africa, UK = United Kingdom, USA = United States of America.

| **Website** | **Number of ads selling individuals under scientific names** | **Number of**  **valid species in the ads** | **Number of ads selling individuals under ‘trade names’** | **Number of ‘trade names’ in the ads** | **Number of valid Genera in the ads** |
| --- | --- | --- | --- | --- | --- |
| CAN | 158 | 135 | 43 | 36 | 62 |
| GER | 102 | 42 | 22 | 8 | 28 |
| POL1 | 191 | 114 | 57 | 43 | 62 |
| POL2 | 81 | 63 | 12 | 8 | 34 |
| SA | 63 | 37 | 5 | 5 | 23 |
| UK | 65 | 51 | 13 | 10 | 35 |
| USA1 | 67 | 58 | 12 | 11 | 36 |
| USA2 | 86 | 58 | 0 | 0 | 35 |

Table S6.2. Number of advertised valid species (n= 217) shared among the websites.

|  | **CAN** | **GER** | **POL1** | **POL2** | **SA** | **UK** | **USA1** | **USA2** |
| --- | --- | --- | --- | --- | --- | --- | --- | --- |
| **CAN** | 0 |  |  |  |  |  |  |  |
| **GER** | 28 | 0 |  |  |  |  |  |  |
| **POL1** | 62 | 30 | 0 |  |  |  |  |  |
| **POL2** | 32 | 13 | 30 | 0 |  |  |  |  |
| **SA** | 22 | 14 | 24 | 14 | 0 |  |  |  |
| **UK** | 31 | 23 | 41 | 18 | 15 | 0 |  |  |
| **USA1** | 39 | 13 | 38 | 15 | 11 | 15 | 0 |  |
| **USA2** | 32 | 18 | 35 | 14 | 12 | 19 | 21 | 0 |
| **Website TOTAL** | 135 | 42 | 114 | 44 | 37 | 51 | 58 | 58 |

**APPENDIX S7 -** *Marginal Means Plots for Categorical Variables*

Marginal means plots for all categorical variables included in (1) species-level and (2) advertisement-level final SEMs.

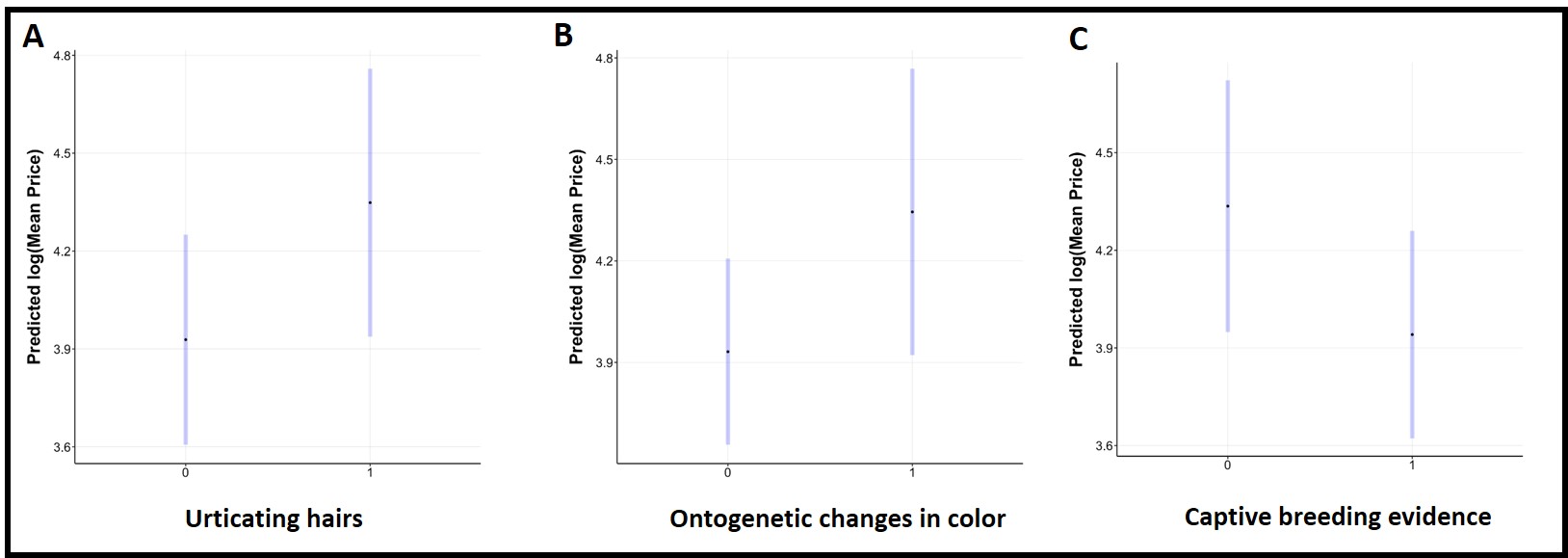

Figure S7.1: marginal means predictions of natural log(mean price) for species **A.** with (1) and without (0) urticating hairs. **B**. with (1) and without (0) ontogenetic changes in color. **C.** with (1) and without (0) captive breeding evidence. 95% confidence intervals are displayed in blue.

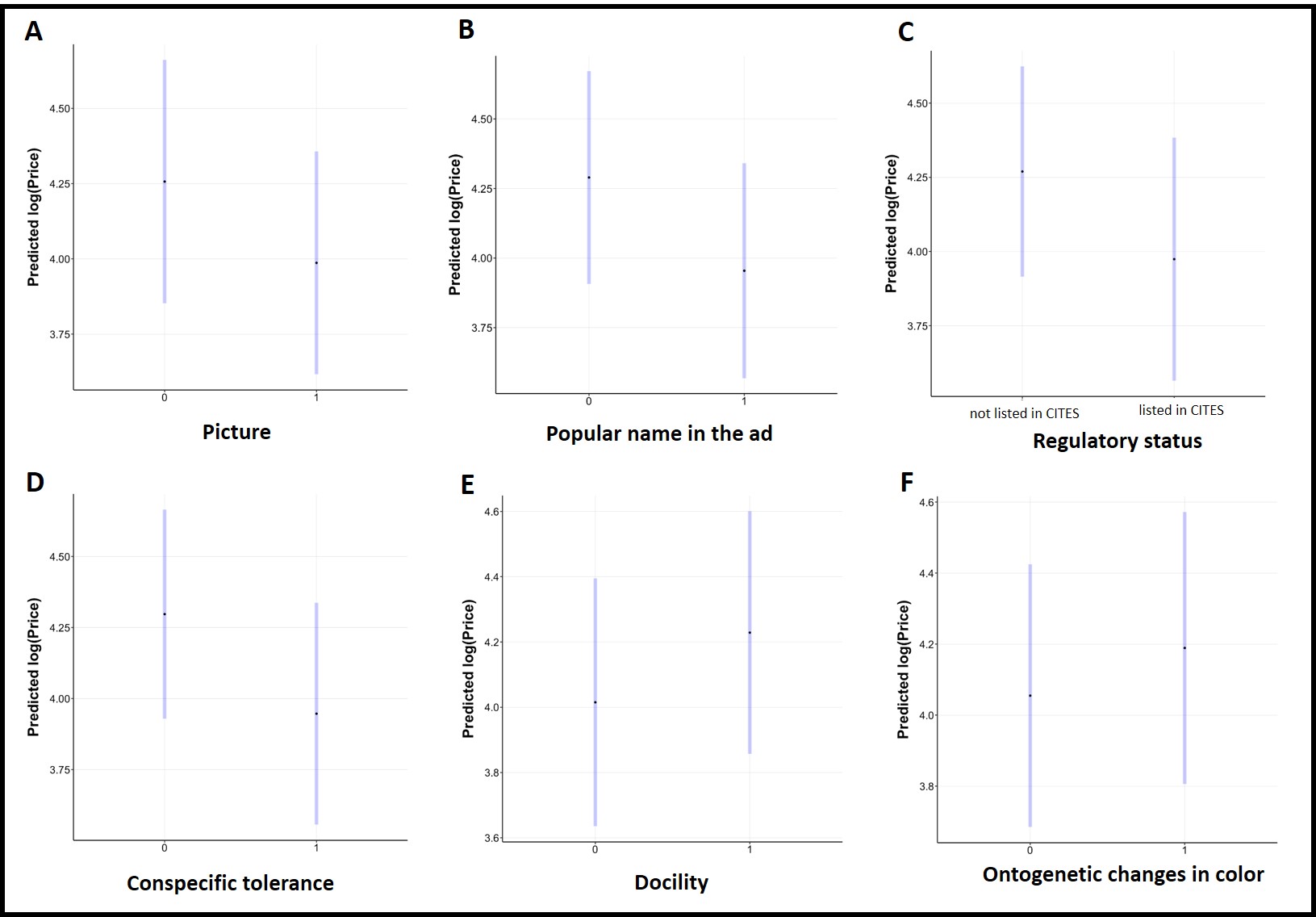

Figure S7.2: marginal means predictions of natural log(price) for advertisements that **A.** do (1) and do not (0) display pictures of the advertised animal. **B**. do (1) and do not (0) display common/popular names of the advertised animal. **C**. display species that are (1) and are not (0) listed in CITES. **D**. display species that have (1) and do not have (0) co-specific tolerance. **E.** display species that have (1) and do not have (0) docile behavior. **F**. have (1) and do not have (0) ontogenetic changes in color. 95% confidence intervals are displayed in blue.

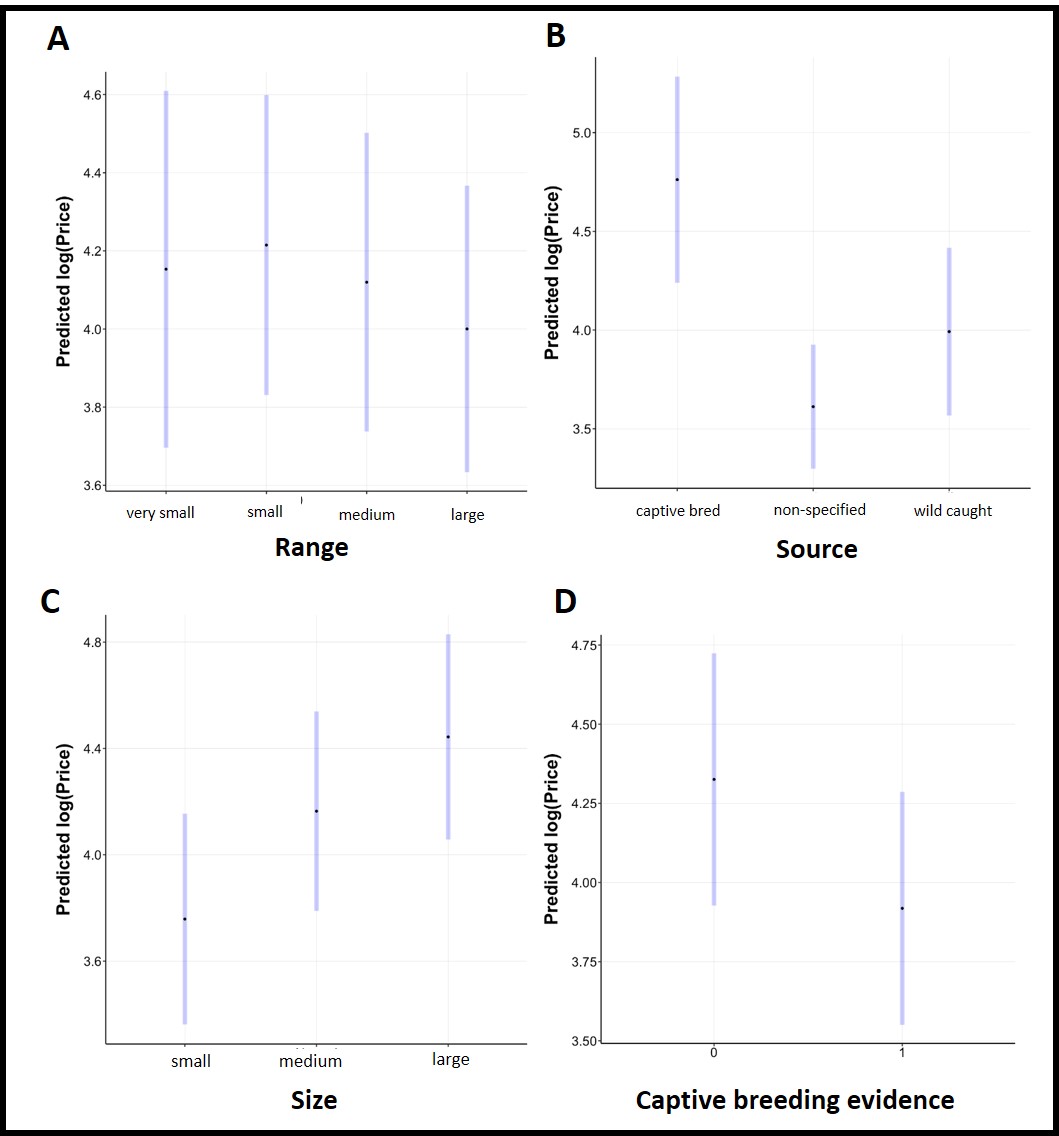

Figure S7.3: marginal means predictions of natural log(price) for advertisements of species **A.** with different range size categories. **B**. with captive bred, wild-caught or non-specified source. **C.** in different size categories. **D.** with (1) and without (0) evidence of captive breeding. 95% confidence intervals are displayed in blue.

**APPENDIX S8** - *SEM analyses: Final Paths*

*Final Paths*

Table S8.1. Final SEM path, standard errors and significance values for species traits. Covariation between variables is indicated with ‘~’. P-values <0.0005 are denoted with ‘***’, P-values <0.001 are denoted with ‘**’ and P-values <0.05 are denoted with ‘*’.Abundance Model *R* ^2^ = 0.56. Average Price Model *R* ^2^ = 0.30

| **Response** | **Predictor** | **Estimate** | **Standard Error** | **Degrees of Freedom** | **Critical Value** | **P-value** | **Standard**  **Estimate** |
| --- | --- | --- | --- | --- | --- | --- | --- |
| Abundance | Global interest | 0.8686 | 0.0735 | 194 | 11.8185 | 0 *** | - |
| Abundance | Growth rate | - 0.1806 | 0.0767 | 194 | - 2.3538 | 0.0186 * | - |
| Abundance | Hairiness score | 0.2382 | 0.0717 | 194 | 3.3207 | 0.0009 *** | - |
| Abundance | Tolerance to conspecifics | - | - | 1 | 0.0078 | 0.9297 | - |
|  | tolerant | 1.5318 | 0.2069 | Inf | 7.4047 | 0 *** | - |
|  | not tolerant | 1.5521 | 0.0724 | Inf | 21.4336 | 0 *** | - |
| Abundance | Docility | - | - | 1 | 11.5623 | 0.0008 *** | - |
|  | docile | 1.2744 | 0.1447 | Inf | 8.8047 | 0 *** | - |
|  | non docile | 1.8095 | 0.1217 | Inf | 14.8672 | 0 *** | - |
| Abundance | Last publication’s authorship | - | - | 1 | 7.0408 | 0.0086 ** | - |
|  | non professional | 1.3531 | 0.1387 | Inf | 9.7534 | 0 *** | - |
|  | professional | 1.7308 | 0.12 | Inf | 14.4286 | 0 *** |  |
| Average Price | Growth rate | 0.0087 | 0.0682 | 192 | 0.1276 | 0.8986 | - |
| Average Price | Year of description (log) | - 0.1137 | 0.0467 | 192 | - 2.4369 | 0.0157 * | - |
| Average Price | Urticating hairs | - | - | 1 | 5.9206 | 0.0159 * | - |
|  | absent | 3.9284 | 0.1633 | 192 | 24.0622 | 0 *** | - |
|  | present | 4.3484 | 0.2082 | 192 | 20.884 | 0 *** | - |
| Average Price | Ontogenetic changes in color | - | - | 1 | 8.2395 | 0.0046 ** | - |
|  | absent | 3.9318 | 0.1394 | 192 | 28.2106 | 0 *** | - |
|  | present | 4.345 | 0.2146 | 192 | 20.2484 | 0 *** | - |
| Average Price | Evidence of captive breeding | - | - | 1 | 8.241 | 0.0046 ** | - |
|  | present | 3.9411 | 0.1616 | 192 | 24.3804 | 0 *** | - |
|  | absent | 4.3357 | 0.196 | 192 | 22.1163 | 0 *** | - |
| Average Price | Size category | - | - | 2 | 9.3882 | 0.0001 *** | - |
|  | small | 3.8915 | 0.2151 | 192 | 18.0952 | 0 *** | - |
|  | medium | 3.9559 | 0.1464 | 192 | 27.0193 | 0 *** | - |
|  | large | 4.5679 | 0.2079 | 192 | 21.9711 | 0 *** | - |
| Average Price | Distribution category | - | - | 5 | 6.8498 | 0 *** | - |
|  | Neotropical | 3.4599 | 0.0987 | 192 | 35.0573 | 0 *** | - |
|  | Afrotropical | 3.9723 | 0.2275 | 192 | 17.4612 | 0 *** | - |
|  | Indo-Malayan | 4.0332 | 0.1949 | 192 | 20.6886 | 0 *** | - |
|  | Palearctic | 4.3693 | 0.5124 | 192 | 8.5268 | 0 *** | - |
|  | Nearctic | 4.481 | 0.2167 | 192 | 20.6808 | 0 *** | - |
|  | Australasian | 4.5147 | 0.4377 | 192 | 10.3135 | 0 *** | - |
| Urticating hairs | ~~Distribution | 0.7016 | - | 224 | 14.7371 | 0 *** | 0.7016 |
| Docility | ~~Urticating Hairs | 0.5319 | - | 211 | 9.1241 | 0 *** | 0.5319 |
| Docility | ~~Distribution | 0.3659 | - | 211 | 5.7109 | 0 *** | 0.3659 |

Tab. S8.2 - Final SEM path, standard errors and significance values for Advertisement trait model, variable response “price” (log transformed) SEM. Covariation between variables is indicated with ‘~’. P-values <0.0005 are denoted with ‘***’, P-values <0.001 are denoted with ‘**’ and P-values <0.05 are denoted with ‘*’. Country codes for the web stores: CAN = Canada, GER = Germany, POL = Poland, SA = South Africa, UK = United Kingdom, USA = United States of America. *R* ^2^ = 0.59.

| **Response** | **Predictor** | **Estimate** | **Standard Error** | **Degrees of Freedom** | **Critical Value** | **P-value** | **Standard**  **Estimate** |
| --- | --- | --- | --- | --- | --- | --- | --- |
| Price (log transformed) | Hairiness score | -0.0907 | 0.0312 | 635 | -2.91 | 0.0037** | - |
| Price (log transformed) | Color score | 0.093 | 0.0319 | 635 | 2.9183 | 0.0036** | - |
| Price (log transformed) | Abundance | -0.0104 | 0.0024 | 635 | -4.3661 | 0*** | - |
| Price (log transformed) | Year of description | -0.1049 | 0.0271 | 635 | -3.8735 | 0.0001*** | - |
| Price (log transformed) | Growth rate | -0.0144 | 0.0353 | 635 | -0.4095 | 0.6823 | - |
| Price (log transformed) | Range category | - | - | 3 | 3.093 | 0.0265* | - |
|  | large | 4.0002 | 0.1866 | 635 | 21.4362 | 0*** | - |
|  | medium | 4.1198 | 0.1947 | 635 | 21.1587 | 0*** | - |
|  | very small | 4.1531 | 0.2325 | 635 | 17.8632 | 0*** | - |
|  | small | 4.2148 | 0.1955 | 635 | 21.5631 | 0*** | - |
| Price (log transformed) | Regulatory status | - | - | 1 | 8.6856 | 0.0033** | - |
|  | present | 3.9743 | 0.2086 | 635 | 19.0547 | 0*** | - |
|  | absent | 4.2697 | 0.1804 | 635 | 23.6653 | 0*** | - |
| Price (log transformed) | Ontogenetic changes in color: | - | - | 1 | 3.6659 | 0.056 | - |
|  | absent | 4.055 | 0.1883 | 635 | 21.5329 | 0*** | - |
|  | present | 4.189 | 0.195 | 635 | 21.4831 | 0*** | - |
| Price (log transformed) | Docility | - | - | 1 | 10.5119 | 0.0012** | - |
|  | non docile | 4.0152 | 0.1932 | 635 | 20.7795 | 0*** | - |
|  | docile | 4.2288 | 0.1894 | 635 | 22.3301 | 0*** | - |
| Price (log transformed) | Tolerance to conspecifics | - | - | 1 | 17.5733 | 0*** | - |
|  | tolerant | 3.9469 | 0.1986 | 635 | 19.8702 | 0*** | - |
|  | not tolerant | 4.2971 | 0.1873 | 635 | 22.9469 | 0*** | - |
| Price (log transformed) | Evidence of captive breeding | - | - | 1 | 15.8547 | 0.0001*** | - |
|  | present | 3.9184 | 0.1874 | 635 | 20.9113 | 0*** | - |
|  | absent | 4.3256 | 0.2028 | 635 | 21.324 | 0*** | - |
| Price (log transformed) | Size category | - | - | 2 | 19.3824 | 0*** | - |
|  | small | 3.7584 | 0.2019 | 635 | 18.6195 | 0*** | - |
|  | medium | 4.1642 | 0.1908 | 635 | 21.8194 | 0*** | - |
|  | large | 4.4435 | 0.1965 | 635 | 22.6139 | 0*** | - |
| Price (log transformed) | Source | - | - | 2 | 13.859 | 0*** | - |
|  | not specified | 3.612 | 0.1599 | 635 | 22.59 | 0*** | - |
|  | wild caught | 3.9922 | 0.2161 | 635 | 18.4706 | 0*** | - |
|  | captive bred | 4.7617 | 0.2658 | 635 | 17.9148 | 0*** | - |
| Price (log transformed) | Picture | - | - | 1 | 5.319 | 0.0214* | - |
|  | present | 3.987 | 0.1885 | 635 | 21.1553 | 0*** | - |
|  | absent | 4.257 | 0.2058 | 635 | 20.6817 | 0*** | - |
| Price (log transformed) | Sex | - | - | 1 | 69.1641 | 0*** | - |
|  | non female | 3.7975 | 0.1905 | 635 | 19.9332 | 0*** | - |
|  | female or couple | 4.4465 | 0.1944 | 635 | 22.8753 | 0*** | - |
| Price (log transformed) | Life stage | - | - | 4 | 19.7312 | 0*** | - |
|  | spiderling | 3.595 | 0.1419 | 635 | 25.3405 | 0*** | - |
|  | juvenile | 4.1068 | 0.1292 | 635 | 31.7879 | 0*** | - |
|  | not specified | 4.1237 | 0.6756 | 635 | 6.1038 | 0*** | - |
|  | adult | 4.3791 | 0.1597 | 635 | 27.4154 | 0*** | - |
|  | subadult | 4.4054 | 0.2139 | 635 | 20.5932 | 0*** | - |
| Price (log transformed) | Website | - | - | 7 | 40.0391 | 0*** | - |
|  | POL2 | 3.4056 | 0.2304 | 635 | 14.7807 | 0*** | - |
|  | POL1 | 3.7456 | 0.2104 | 635 | 17.8055 | 0*** | - |
|  | USA2 | 3.928 | 0.1969 | 635 | 19.945 | 0*** | - |
|  | SA | 3.9499 | 0.2169 | 635 | 18.2144 | 0*** | - |
|  | UK | 4.116 | 0.2233 | 635 | 18.431 | 0*** | - |
|  | GER | 4.2884 | 0.2337 | 635 | 18.3528 | 0*** | - |
|  | CAN | 4.4266 | 0.2167 | 635 | 20.4254 | 0*** | - |
|  | USA1 | 5.1158 | 0.2271 | 635 | 22.5251 | 0*** | - |
| Price (log transformed) | Popular name in the ad | - | - | 1 | 10.1128 | 0.0015** | - |
|  | present | 3.9546 | 0.1967 | 635 | 20.1006 | 0*** | - |
|  | absent | 4.2894 | 0.1946 | 635 | 22.0424 | 0*** | - |
| Urticating hairs | ~~Distribution | 0.6773 | - | 707 | 24.4767 | 0*** | 0.6773 |
| Docility | ~~Urticating hairs | 0.4944 | - | 686 | 14.8989 | 0*** | 0.4944 |
| Docility | ~~Distribution | 0.2808 | - | 686 | 7.6618 | 0*** | 0.2808 |

Tab. S8.3 - Final SEM path, standard errors and significance values for Advertisement trait model, Mixed Model, variable response “price” (log transformed) SEM. Covariation between variables is indicated with ‘~’. P-values <0.0005 are denoted with ‘***’, P-values <0.001 are denoted with ‘**’ and P-values <0.05 are denoted with ‘*’. For the predictors marked with ^a^ , the response variable is “urticating hairs”, and for ^b^ the response variable is “docility”. *R* ^2^ =.0.57.

| **Response** | **Predictor** | **Estimate** | **Standard Error** | **Degrees of Freedom** | **Critical Value** | **P-value** | **Standard**  **Estimate** |
| --- | --- | --- | --- | --- | --- | --- | --- |
| Price (log transformed) | Hairiness score | -0.0865 | 0.0351 | 577.2199 | -2.4645 | 0.014* | - |
| Price (log transformed) | Color score | 0.0996 | 0.0339 | 577.377 | 2.9412 | 0.0034** | - |
| Price (log transformed) | Abundance | -0.0102 | 0.0026 | 577.4793 | -3.9939 | 0.0001*** | - |
| Price (log transformed) | Year of description (log) | -0.1122 | 0.0292 | 577.5164 | -3.841 | 0.0001*** | - |
| Price (log transformed) | Size advertised | -6.00E-04 | 9.00E-04 | 577.4126 | -0.6912 | 0.4897 | - |
| Price (log transformed) | Growth rate | -0.0725 | 0.0433 | 577.5389 | -1.6768 | 0.0941 | - |
| Price (log transformed) | Picture | - | - | 1 | 7.5998 | 0.006** | - |
|  | present | 4.0193 | 0.2638 | 13.8926 | 15.2338 | 0*** | - |
|  | absent | 4.2919 | 0.2728 | 15.8018 | 15.7316 | 0*** | - |
| Price (log transformed) | Regulatory status | - | - | 1 | 7.1081 | 0.0079** | - |
|  | present | 4.011 | 0.2809 | 17.7689 | 14.2807 | 0*** | - |
|  | absent | 4.3002 | 0.2572 | 12.5346 | 16.72 | 0*** | - |
| Price (log transformed) | Urticating hairs: | - | - | 1 | 3.5217 | 0.0611 | - |
|  | absent | 4.0495 | 0.26 | 13.1156 | 15.5732 | 0*** | - |
|  | present | 4.2617 | 0.2792 | 17.3117 | 15.2658 | 0*** | - |
| Price (log transformed) | Evidence of captive breeding: | - | - | 1 | 19.7216 | 0*** | - |
|  | present | 3.9175 | 0.2602 | 13.1279 | 15.054 | 0*** | - |
|  | absent | 4.3937 | 0.2778 | 17.0272 | 15.8149 | 0*** | - |
| Price (log transformed) | Tolerance to conspecifics | - | - | 1 | 19.2361 | 0*** | - |
|  | present | 3.9131 | 0.2708 | 15.3812 | 14.4481 | 0*** | - |
|  | absent | 4.3981 | 0.2682 | 14.8045 | 16.4009 | 0*** | - |
| Price (log transformed) | Docility | - | - | 1 | 11.7255 | 0.0007*** | - |
|  | absent | 4.0276 | 0.266 | 14.3275 | 15.1415 | 0*** | - |
|  | present | 4.2836 | 0.2668 | 14.5067 | 16.0545 | 0*** | - |
| Price (log transformed) | Size category | - | - | 2 | 14.1696 | 0*** | - |
|  | small | 3.8798 | 0.2851 | 18.8126 | 13.6085 | 0*** | - |
|  | medium | 4.0795 | 0.2565 | 12.4053 | 15.9054 | 0*** | - |
|  | large | 4.5075 | 0.2748 | 16.3165 | 16.4031 | 0*** | - |
| Price (log transformed) | Distribution category | - | - | 5 | 3.8264 | 0.002** | - |
|  | Neotropical | 3.5945 | 0.235 | 8.7712 | 15.295 | 0*** | - |
|  | Nearctic | 3.909 | 0.2805 | 17.7103 | 13.9346 | 0*** | - |
|  | Afrotropical | 4.0512 | 0.2683 | 14.8464 | 15.0984 | 0*** | - |
|  | Indo-Malayan | 4.0756 | 0.2673 | 14.5872 | 15.2484 | 0*** | - |
|  | Australasian | 4.5297 | 0.383 | 58.4839 | 11.8277 | 0*** | - |
|  | Palearctic | 4.7736 | 0.5458 | 191.684 | 8.7456 | 0*** | - |
| Price (log transformed) | Sex advertised | - | - | 1 | 48.3211 | 0*** | - |
|  | non-female | 3.8515 | 0.2681 | 14.7225 | 14.3671 | 0*** | - |
|  | female or couple | 4.4597 | 0.2667 | 14.4989 | 16.7236 | 0*** | - |
| Price (log transformed) | Life stage | - | - | 3 | 23.1591 | 0*** | - |
|  | spiderling | 3.6413 | 0.2562 | 12.3779 | 14.2125 | 0*** | - |
|  | juvenile | 4.1495 | 0.2531 | 11.7839 | 16.3953 | 0*** | - |
|  | subadult | 4.2489 | 0.3811 | 57.3737 | 11.1492 | 0*** | - |
|  | adult | 4.5827 | 0.2893 | 19.8192 | 15.843 | 0*** | - |
| Price (log transformed) | Ontogenetic changes in color | - | - | 1 | 15.4064 | 0.0001*** | - |
|  | absent | 3.9694 | 0.2558 | 12.2651 | 15.5173 | 0*** | - |
|  | present | 4.3418 | 0.2797 | 17.4748 | 15.5247 | 0*** | - |
| Urticating hairs | ~~Distribution ^a^ | 0.6773 | - | 707 | 24.4767 | 0*** | 0.6773 |
| Docility | ~~Urticating Hairs ^b^ | 0.4944 | - | 686 | 14.8989 | 0*** | 0.4944 |
| Docility | ~~Distribution ^b^ | 0.2808 | - | 686 | 7.6618 | 0*** | 0.2808 |

**APPENDIX S9 -** *Relative Variable Importance Scores*

Table S9.1: Relative variable importance scores (standardized to sum to 1) for all variables included as explanatory variables in (**A**) the negative binomial GLM component and (**B**) the linear regression component of the species-level final SEM.

Table S9.1A

| **Variable** | **Relative Variable Importance** |
| --- | --- |
| Global interest | 0.696 |
| Hairiness score | 0.169 |
| Docility | 0.0641 |
| Last publication’s authorship | 0.0360 |
| Growth rate | 0.0270 |
| Tolerance to conspecifics | 0.00880 |

Table S9.1B

| **Variable** | **Relative Variable Importance** |
| --- | --- |
| Distribution category | 0.437 |
| Size category | 0.223 |
| Year of description (log) | 0.112 |
| Evidence of captive breeding | 0.0980 |
| Ontogenetic changes in color | 0.0580 |
| Presence of urticating hairs | 0.0564 |
| Growth rate | 0.0133 |

Table S9.2: Relative variable importance scores (standardized to sum to 1) for all variables included as explanatory variables in the linear regression component of the advertisement-level final SEM.

| **Variable** | **Relative Variable Importance** |
| --- | --- |
| Website | 0.359 |
| Life stage advertised | 0.136 |
| Sex | 0.114 |
| Source | 0.087 |
| Size category | 0.053 |
| Abundance | 0.049 |
| Picture | 0.042 |
| Year of description (log) | 0.035 |
| Evidence of captive breeding | 0.025 |
| Regulatory status | 0.021 |
| Hairiness score | 0.019 |
| Range | 0.016 |
| Docility | 0.013 |
| Popular name in the ad | 0.011 |
| Tolerance to conspecifics | 0.010 |
| Ontogenetic changes in color | 0.005 |
| Color score | 0.004 |
| Growth rate | 0.002 |

**APPENDIX S10**- *Tukey’s pairwise comparison test*

Tukey’s pairwise comparison tests for all categorical variables (with greater than two levels) included in (**A**) species-level and (**B**) advertisement-level final SEMs.

Table S10.A1: Tukey’s pairwise comparisons testing differences in species market abundance between different distribution categories.

| contrast | estimate | SE | df | t.ratio | p.value |
| --- | --- | --- | --- | --- | --- |
| Afrotropical-Australasian | -0.5424 | 0.417 | 192 | -1.301 | 0.7840 |
| Afrotropical-Indo-Malayan | -0.0609 | 0.193 | 192 | -0.315 | 0.9996 |
| Afrotropical-Nearctic | -0.5087 | 0.318 | 192 | -1.598 | 0.6009 |
| Afrotropical-Neotropical | 0.5125 | 0.234 | 192 | 2.188 | 0.2483 |
| Afrotropical-Palearctic | -0.3969 | 0.502 | 192 | -0.791 | 0.9688 |
| Australasian-Australasian | 0.4815 | 0.415 | 192 | 1.159 | 0.8556 |
| Australasian-Nearctic | 0.0337 | 0.491 | 192 | 0.069 | 1.0000 |
| Australasian-Neotropical | 1.0549 | 0.44 | 192 | 2.398 | 0.1620 |
| Australasian-Palearctic- | 0.1455 | 0.622 | 192 | 0.234 | 0.9999 |
| Indo-Malayan-Nearctic | -0.4478 | 0.297 | 192 | -1.508 | 0.6593 |
| Indo-Malayan-Neotropical | 0.5734 | 0.216 | 192 | 2.659 | 0.0885 |
| Indo-Malayan-Palearctic | -0.3361 | 0.497 | 192 | -0.677 | 0.9843 |
| Nearctic-Neotropical | 1.0212 | 0.209 | 192 | 4.882 | <.0001 |
| Nearctic-Palearctic- | 0.1118 | 0.555 | 192 | 0.202 | 1.0000 |
| Neotropical-Palearctic | -0.9094 | 0.516 | 192 | -1.763 | 0.4924 |

Table S10.B1: Tukey’s pairwise comparisons testing differences in species market abundance between different size categories.

| contrast | estimate | SE | df | t.ratio | p.value |
| --- | --- | --- | --- | --- | --- |
| small-medium | -0.406 | 0.0963 | 635 | -4.215 | 0.0001 |
| small-large | -0.685 | 0.1102 | 635 | -6.218 | <.0001 |
| medium-large | -0.279 | 0.0792 | 635 | -3.527 | 0.0013 |

Table S10.B2: Tukey’s pairwise comparisons testing differences in mean log(price) between different range categories.

| contrast | estimate | SE | df | t.ratio | p.value |
| --- | --- | --- | --- | --- | --- |
| very small-small | -0.062 | 0.1617 | 635 | -0.382 | 0.9811 |
| very small-medium | 0.0333 | 0.1557 | 635 | 0.214 | 0.9965 |
| very small-large | 0.1529 | 0.1527 | 635 | 1.001 | 0.7488 |
| small-medium | 0.095 | 0.0886 | 635 | 1.073 | 0.7063 |
| small-large | 0.2146 | 0.0792 | 635 | 2.709 | 0.0349 |
| medium-large | 0.1195 | 0.0624 | 635 | 1.915 | 0.2227 |

Table S10.B3: Tukey’s pairwise comparisons testing differences in mean log(price) between different size categories.

| contrast | estimate | SE | df | t.ratio | p.value |
| --- | --- | --- | --- | --- | --- |
| small-medium | -0.406 | 0.0963 | 635 | -4.215 | 0.0001 |
| small-large | -0.685 | 0.1102 | 635 | -6.218 | <.0001 |
| medium-large | -0.279 | 0.0792 | 635 | -3.527 | 0.0013 |

Table S10.B4: Tukey’s pairwise comparisons testing differences in mean log(price) between different source categories. cb = captive bred, ns = not specified, wc = wild caught.

| contrast | estimate | SE | df | t.ratio | p.value |
| --- | --- | --- | --- | --- | --- |
| cb-ns | 1.15 | 0.222 | 635 | 5.17 | <.0001 |
| cb-wc | 0.77 | 0.181 | 635 | 4.262 | 0.0001 |
| ns-wc | -0.38 | 0.165 | 635 | -2.309 | 0.0552 |

Table S10.B5: Tukey’s pairwise comparisons testing differences in mean log(price) between different life stage categories. NS = not specified.

| contrast | estimate | SE | df | t.ratio | p.value |
| --- | --- | --- | --- | --- | --- |
| adult-juvenile | 0.2722 | 0.1106 | 635 | 2.461 | 0.1009 |
| adult-NS | 0.2553 | 0.6764 | 635 | 0.377 | 0.9957 |
| adult-spiderling | 0.7841 | 0.1206 | 635 | 6.503 | <.0001 |
| adult-subadult | -0.0264 | 0.1705 | 635 | -0.155 | 0.9999 |
| juvenile-NS | -0.0169 | 0.6687 | 635 | -0.025 | 1.0000 |
| juvenile-spiderling | 0.5118 | 0.0643 | 635 | 7.954 | <.0001 |
| juvenile-subadult | -0.2986 | 0.1543 | 635 | -1.936 | 0.2994 |
| NS-spiderling | 0.5287 | 0.6711 | 635 | 0.788 | 0.9341 |
| NS-subadult | -0.2817 | 0.6851 | 635 | -0.411 | 0.9940 |
| spiderling-subadult | -0.8104 | 0.1631 | 635 | -4.968 | <.0001 |

Table S10.B6: Tukey’s pairwise comparisons testing differences in mean log(price) between different websites. Country codes for the web stores: CAN = Canada, GER = Germany, POL = Poland, SA = South Africa, UK = United Kingdom, USA = United States of America.

| contrast | estimate | SE | df | t.ratio | p.value |
| --- | --- | --- | --- | --- | --- |
| CAN-GER | 1.1659 | 0.119 | 635 | 9.768 | <.0001 |
| CAN-POL1 | 1.7102 | 0.154 | 635 | 11.094 | <.0001 |
| CAN-POL2 | 0.8274 | 0.182 | 635 | 4.551 | 0.0002 |
| CAN-SA | 1.3702 | 0.108 | 635 | 12.695 | <.0001 |
| CAN-UK | 0.6892 | 0.14 | 635 | 4.931 | <.0001 |
| CAN-USA1 | 1.1878 | 0.237 | 635 | 5.002 | <.0001 |
| CAN-USA2 | 0.9998 | 0.119 | 635 | 8.398 | <.0001 |
| GER-POL1 | 0.5442 | 0.154 | 635 | 3.526 | 0.0106 |
| GER-POL2 | -0.3385 | 0.178 | 635 | -1.904 | 0.5487 |
| GER-SA | 0.2043 | 0.102 | 635 | 2.009 | 0.4768 |
| GER-UK | -0.4767 | 0.142 | 635 | -3.364 | 0.0184 |
| GER-USA1 | 0.0219 | 0.228 | 635 | 0.096 | 1.0000 |
| GER-USA2 | -0.1661 | 0.117 | 635 | -1.423 | 0.8464 |
| POL1-POL2 | -0.8828 | 0.144 | 635 | -6.135 | <.0001 |
| POL1-SA | -0.34 | 0.119 | 635 | -2.864 | 0.0818 |
| POL1-UK | -1.021 | 0.1 | 635 | -10.201 | <.0001 |
| POL1-USA1 | -0.5224 | 0.262 | 635 | -1.997 | 0.4847 |
| POL1-USA2 | -0.7103 | 0.155 | 635 | -4.594 | 0.0001 |
| POL2-SA | 0.5428 | 0.148 | 635 | 3.67 | 0.0064 |
| POL2-UK | -0.1382 | 0.135 | 635 | -1.02 | 0.9714 |
| POL2-USA1 | 0.3604 | 0.273 | 635 | 1.318 | 0.8921 |
| POL2-USA2 | 0.1724 | 0.182 | 635 | 0.948 | 0.9811 |
| SA-UK | -0.681 | 0.103 | 635 | -6.599 | <.0001 |
| SA-USA1 | -0.1824 | 0.225 | 635 | -0.81 | 0.9925 |
| SA-USA2 | -0.3704 | 0.107 | 635 | -3.458 | 0.0135 |
| UK-USA1 | 0.4986 | 0.248 | 635 | 2.014 | 0.4732 |
| UK-USA2 | 0.3106 | 0.141 | 635 | 2.21 | 0.3469 |
| USA1-USA2 | -0.1879 | 0.23 | 635 | -0.815 | 0.9922 |
